# Monitoring the translation initiation process in prokaryotes via large-scale examination using optimized SSU-seq technique

**DOI:** 10.64898/2026.10.04.756509

**Authors:** Larissa Fine, Alon Diament, Rachel Cohen-Kupiec, Tamir Tuller

**Affiliations:** Department of Biomedical Engineering, Tel Aviv University, Tel Aviv 69978, Israel

## Abstract

Translation initiation is a critical stage of protein synthesis in prokaryotes, where ribosomes assemble on mRNA to begin decoding genetic information. The interactions between ribosomal RNA (rRNA) and mRNA significantly influence this process, with features like ribosomal binding site (RBS) affecting ribosomal dynamics. However, direct experimental validation of these types of interaction is yet to be provided.

In this study, we developed an optimized small sub-unit sequencing (SSU-seq) protocol for *Escherichia coli*, enabling the detailed examination of ribosomal complexes and subunits dynamics. Unlike Ribo-seq, which primarily maps fully assembled ribosomes, SSU-seq captures both ribosomal subunits and intact ribosomes, offering a comprehensive view of translation initiation and elongation events. By integrating experimental data with predictive models, our study advances the understanding of translation regulation in prokaryotes and underscores the utility of SSU-seq in decoding the complexities of ribosome-mRNA interactions. Our analysis reveals distinct patterns of small subunit distribution across the entire mRNAs with higher density near positions with stronger rRNA-mRNA hybridization. The results may suggest that rRNA-mRNA hybridization positions along the entire transcript facilitate efficient start codon recognition by maintaining proximity between the small subunit and the mRNA.

## INTRODUCTION

In prokaryotes, translation initiation is a tightly regulated and often rate-limiting step that determines ribosome positioning and overall protein synthesis efficiency. Initiation involves recognition of the ribosome-binding site by the 30S ribosomal subunit, mediated by base-pairing between the mRNA Shine–Dalgarno sequence and the complementary anti–Shine– Dalgarno region of the 16S rRNA, which aligns the start codon in the ribosomal P site. Beyond canonical Shine–Dalgarno interactions, accumulating evidence indicates that additional rRNA–mRNA contacts and mRNA structural features modulate initiation efficiency, start-site selection, and ribosome recruitment dynamics^1–3^.

Building on this broader view of rRNA involvement in translation, a predictive model was developed in prokaryotes to describe small ribosomal subunit hybridization dynamics along mRNA^4^. The model proposed that transient rRNA–mRNA interactions retain free small ribosomal subunits near the transcript before start-codon recognition, increasing initiation probability. It distinguished between strong, intermediate, and weak hybridization states (Figure 1): strong interactions may trap the subunit away from productive start sites, whereas weak interactions allow premature dissociation. In contrast, intermediate hybridization promotes repeated binding and release events that keep the subunit near the transcript until start-codon recognition. However, direct experimental evidence for these interactions in vivo remained lacking.

**Figure 1:**
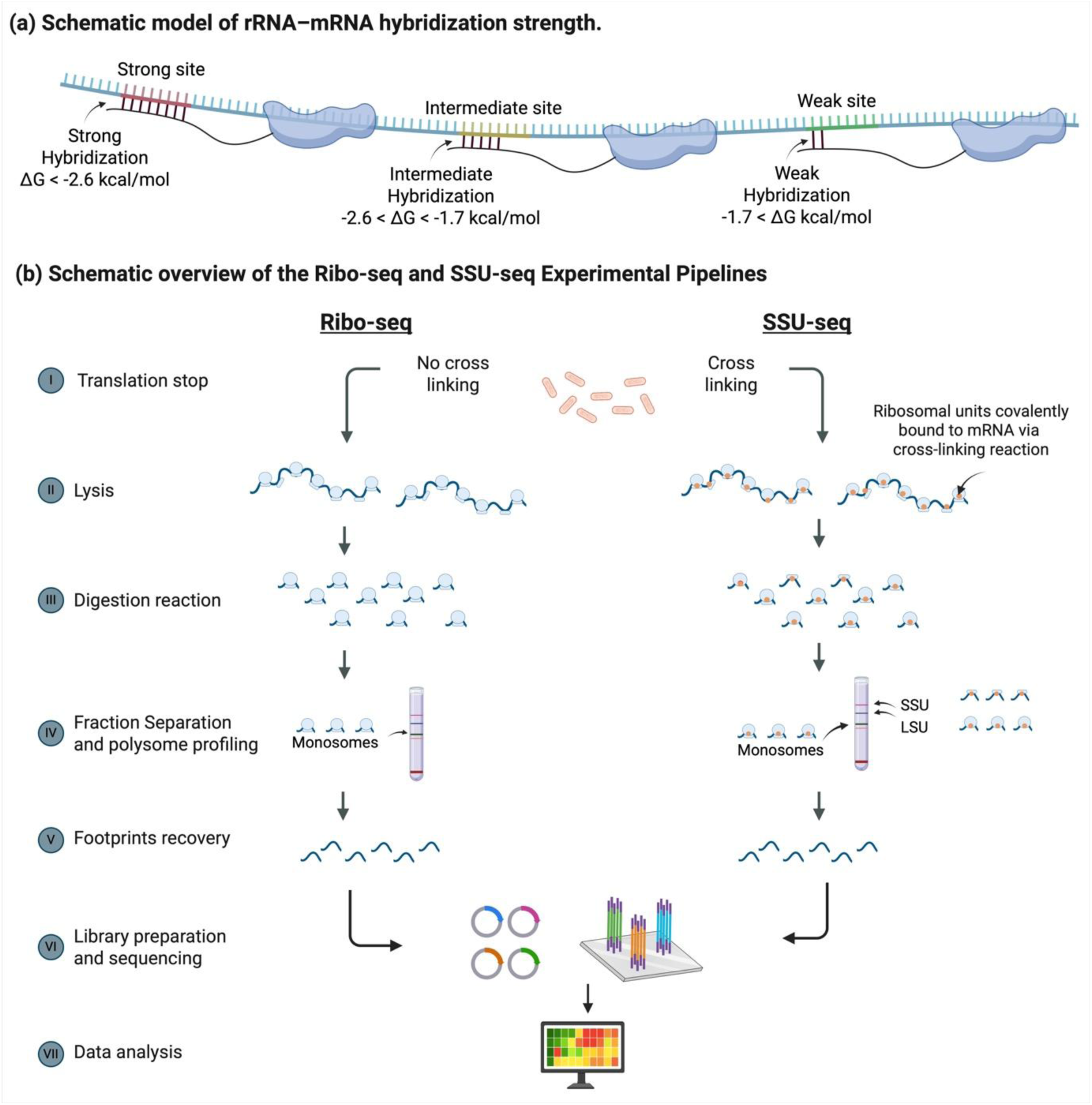
Schematic representation of rRNA–mRNA hybridization strength and overview of the Ribo-seq and SSU-seq experimental pipelines. (a) Schematic model of predicted rRNA– mRNA hybridization interactions, illustrating strong, intermediate, and weak hybridization sites along mRNA. (b) Schematic overview of the Ribo-seq and SSU-seq experimental pipelines. Both workflows include (I) translation arrest, achieved by rapid freezing of cells in liquid nitrogen, with an additional formaldehyde cross-linking step in the SSU-seq protocol; (II) cell lysis using combined chemical lysis and mechanical disruption by cryogenic pulverization; (III) mRNA digestion using micrococcal nuclease (MNase); (IV) polysome profiling and fraction separation by sucrose gradient centrifugation; (V) ribosome-protected footprint recovery by phenol extraction; (VI) library preparation and sequencing; and (VII) downstream data analysis. In contrast to Ribo-seq, SSU-seq preserves ribosomal subunit–mRNA interactions through covalent cross-linking, enabling the recovery of footprints associated with small ribosomal subunits (SSU), large subunits (LSU), and monosomes.

Ribo-seq (Ribosome profiling sequencing) is a high-throughput sequencing technique that enables genome-wide measurement of active translation by capturing ribosome-protected mRNA fragments. By sequencing ∼28–30 nucleotide fragments shielded from nuclease digestion, Ribo-seq provides nucleotide-resolution maps of ribosome occupancy along transcripts and has substantially advanced the study of translational regulation^4^. By design, Ribo-seq reports footprints derived from translating ribosomes and does not interrogate the positioning or interaction states of individual ribosomal subunits during early initiation events. As a result, rRNA–mRNA interactions involving the small ribosomal subunit prior to or during initiation remain outside the scope of conventional Ribo-seq^5–7^.

In this study, we developed and optimized an SSU-seq (small subunit sequencing) protocol for prokaryotes, implementing it in *Escherichia coli* to enable genome-wide interrogation of ribosomal small subunit–mRNA interactions. By adapting the SSU-seq framework to selectively capture SSU-associated footprints, this approach allows direct mapping of small subunit positioning along transcripts, rather than inferring initiation events from assembled ribosomes. Using SSU-seq, we systematically characterize the distribution of the ribosomal small subunit across distinct transcript regions, providing new insights into early translation events and rRNA–mRNA interaction landscapes in bacteria. Together, this work establishes SSU-seq as a dedicated experimental framework for studying bacterial translation initiation and SSU engagement at transcriptome scale.

## MATERIAL AND METHODS

### Experimental Methods

The protocol of SSU-seq was optimized to maximize footprint recovery and concentration, ensuring that the SSU-seq protocol provides an accurate representation of the translational landscape, from initiation to termination. The full protocol is described in the supplementary section, while the main optimizations are described below.

### Medium and cross-linking reaction

*E. coli* cultures were grown to mid-log phase to ensure high ribosomal activity and strong translation signals. To optimize growth conditions and prevent stress-induced gene expression changes, we compared LB (rich, undefined) and MOPS EZ (rich, defined) media. Polysome profiling showed that MOPS EZ yielded higher footprint concentrations (Figure S 1a), making it the preferred medium for subsequent experiments^8–11^.

To preserve ribosome-mRNA interactions at various translation stages, formaldehyde cross-linking was chosen over alternatives like cycloheximide (which only captures elongating ribosomes and may distort translation efficiency measurements^12,13^ and harringtonine (which primarily captures initiation events^14,15^). The efficiency of cross-linking depends on formaldehyde concentration, pH, and other conditions^16–18^.

To optimize these parameters, we simultaneously tested the effects of medium type and cross-linking pH. Figure S 1a shows RNA extracted from *E. coli* cultures grown in LB or MOPS media, subjected to three cross-linking conditions: (1) low pH <5, (2) pH 8, and (3) no cross-linking. In all samples (lanes 2–7), a large band at the bottom represents tRNAs. MOPS samples show two distinct rRNA bands under all pH conditions, while RNA concentration from LB-grown cultures is notably lower, reinforcing the choice of MOPS for SSU-seq. Focusing on MOPS samples, the reference sample without cross-linking (lane 7) exhibited the highest rRNA concentration, while lanes 5 and 6 (low pH and pH 8) showed reduced rRNA bands. OD readings after ethanol precipitation (not shown) were similar, with a slight advantage for alkaline conditions, consistent with improved RNA fixation at higher pH^19,20^. However, inconclusive pH effects suggest that glycine quenching may have been insufficient, potentially leading to over-fixation and reduced Trizol extraction efficiency. We further optimized formaldehyde concentration, testing 0%, 0.5%, 1%, 1.5%, and 2%. Polysome profiles (Figure S 1) revealed an inverse relationship between formaldehyde concentration and monosome levels, with higher concentrations increasing SSU fraction. Based on these results, 0.5% formaldehyde was selected as optimal for capturing SSU, LSU, and monosome fractions. Following cross-linking, cells were flash-frozen in liquid nitrogen to halt all cellular processes.

### Cell lysis

We optimized the *E. coli* cell lysis protocol by combining mechanical grinding with chemical lysis to enhance disruption efficiency. The optimal cell amount (in grams) for each method was determined. The optimized program includes a 5-minute pre-cooling step, followed by 5 cycles at 20 Hz, each consisting of 2 minutes of grinding and 1 minute of cooling. This regimen consistently yields a fine cell powder.

### Nuclease digestion

Micrococcal Nuclease (MNase) digestion targets exposed mRNA regions not protected by ribosomes, leveraging its endonucleolytic and exonucleolytic activity. Polysome profiling confirms that MNase digestion increases monosome levels (Figure S 1 c,d), indicating polysome cleavage into ribosome-protected fragments. To optimize digestion, we compared reactions at room temperature (RT) and 37°C across at least three independent replicates (Figure S 1e,f). Both conditions yielded similar profiles, but to prevent excessive degradation, the reaction was performed at RT for one hour, ensuring robust and reproducible results. MNase exhibits a substantial sequence preference for AT-rich substrates, cleaving approximately 30-fold faster on the 5′ side of A or T than of G or C^21^; the same bias has been characterised in bacterial ribosome profiling^13,22^ Consequently, read 5′-ends are systematically depleted within the GC-rich predicted Shine-Dalgarno-like motifs themselves, visible as a narrow vertical depletion band at displayed positions ∼+1 to +5 in the Figure 4 metagene heatmaps; this reflects a technical bias of the assay rather than a feature of ribosomal subunit positioning.

### Polysome profiling

Polysome profiling enables the analysis of translation dynamics by fractionating cell lysates on a 10%–50% (w/v) sucrose gradient. The gradient, prepared using a BIOCOMP fractionator, was loaded with digested mRNA bound to ribosomal subunits and subjected to ultracentrifugation in an SW-41 rotor for density-based separation. The resulting polysome profile revealed ribosomal subunit and mRNA complex distribution. Individual fractions were collected and flash-frozen immediately to preserve integrity for further analysis.

### Footprint isolation

Following fractionation, mRNA footprints were separated from ribosomal subunits by reversing the cross-linking reaction. Phenol-chloroform extraction was used to dissociate ribosomal components from mRNA, with heat treatment duration optimized for maximum footprint recovery (data not shown). RNA fragments (15–70 nucleotides) were resolved on a 15% TBE-urea gel, excised (Figure S 2a, red boxes), and recovered using the PAGExt Gel Extraction Kit. The quality and size distribution of the isolated footprints were assessed using the 2100 Bioanalyzer with the Agilent High Sensitivity DNA Kit, ensuring high-quality footprint isolation and library preparation for subsequent analyses (Figure S 2b).

### Library preparation and quality check

libraries were prepared by IMMAGINA using LACEseq kit. The quality of the resulting libraries was assessed on an Agilent 2100 Bioanalyzer system.

### Sequencing

Libraries were sequenced on the NovaSeq™ 6000 instrument.

### Size selection

In eukaryotic SSU-seq, footprint sizes typically range from 15 to 75 nucleotides. In prokaryotes, ribosome-protected mRNA fragments are generally 20–40 nucleotides long, with most footprints clustering around 28–30 nucleotides^23,24^. The broader range in SSU-seq is due to its ability to capture footprints from both the small subunit and the full ribosome, along with associated initiation factors^25^. Since no prior SSU-seq studies have been conducted in prokaryotes, we adopted a similar size range (15–70 nucleotides) as used in eukaryotic studies for our initial experiments. This broader selection enabled a more comprehensive analysis and optimization based on subsequent data. Footprint isolation within this range was performed in collaboration with IMMAGINA BioTechnology.

## Computational Methods

### Read mapping

Transcript sequences were obtained from NCBI for E. coli (NC_000913.3). Sequenced reads were mapped as follows^26^. Transcript 5′ UTR and 3′ UTR boundaries were defined as 50 bp upstream of each annotated start codon and up to 100 bp downstream of each stop codon, trimmed at the midpoint of any intergenic region with the next same-strand gene where applicable but never below a 50-bp minimum length. The 50-bp 5′ UTR window approximates the median 5′ UTR length in E. coli K-12, which is 43–57 nt across genes with an experimentally mapped transcription start site, depending on whether the shortest or longest transcription unit is taken for genes transcribed from several ^27^. We trimmed 3’ adaptors from the reads using Cutadapt^28,29^, UMI barcodes with umi-tools^30^, and utilized Bowtie^31^ to map them to the E.coli transcriptome. In the first phase, we discarded reads that mapped to rRNA and tRNA sequences with Bowtie parameters ‘–n 2 –seedlen 21 –k 1 --norc’. In the second phase, we mapped the remaining reads to the transcriptome with Bowtie parameters ‘–v 2 –a --strata --best --norc –m 200’. We filtered out reads longer than 75nt and shorter than 15nt. Unique alignments were first assigned to the occupancy profiles. For multiple alignments, the best alignments in terms of number of mismatches were kept. Then, multiple aligned reads were distributed between locations according to the distribution of unique reads in the respective surrounding regions. To this end, a 100nt window was used to compute the read count density RCDi (total read counts in the window divided by length, based on unique reads) in vicinity of the M multiple aligned positions in the transcriptome, and the fraction of a read assigned to each position was RCDij=1MRCDj.

### rRNA–mRNA Hybridization Strength

rRNA–mRNA hybridization strength was calculated based on predicted hybridization free energy between short mRNA subsequences and the aSD sequence at the 3′ end of the 16S rRNA, as previously described by Bahiri et al. (2021)^5^. Hybridization energies were computed for all possible 6-nt windows along each transcript using RNAcoFold from the ViennaRNA package. Profiles were available for the 50 nt upstream of each start codon, the coding sequence, and the first 45 nt of each 3′ UTR; all site-based analyses are therefore restricted to these intervals. The 45-nt 3′ UTR interval covers the full 3′ UTR for the majority of genes with an experimentally mapped transcription terminator (median 3′ UTR length 19–49 nt, depending on which edge of the terminator is taken; RegulonDB v12). More negative free energy values correspond to stronger rRNA–mRNA interactions.

Interaction strength thresholds were adopted from the previously published study, which defined interaction categories using an unsupervised optimization framework comparing native genomes to randomized transcriptomes. Based on these thresholds, interaction strengths were classified as (1) **strong interactions:** ΔG < −2.6 kcal/mol. (2) **Intermediate interactions:** −2.6 ≤ ΔG < −1.7 kcal/mol. (3) **Weak interactions:** ΔG ≥ −1.7 kcal/mol. In the 5′ UTR, 51% of intermediate-strength sites lie within 3 nt of a strong-strength site in the same gene, reflecting the sliding-window nature of the hybridization-energy profile rather than distinct motif populations.

These thresholds were applied genome-wide to identify predicted rRNA–mRNA hybridization sites across transcripts.

Per-gene hybridization energy was taken as the strongest predicted value in the canonical Shine–Dalgarno register, 8–15 nt upstream of the start codon. Occupancy was measured in a fixed 16-nt window 15–30 nt upstream of the start codon, where the 5′ ends of initiation footprints fall, and normalized by the read density in the first half of the same gene’s coding sequence, trimmed by 60 nt at each end so that initiation and termination peaks are excluded from the denominator. Because every gene contributes at the same position relative to the start codon, distance to the start codon is held constant by construction. Genes were binned into energy quintiles, and the statistic was the fraction of genes whose occupancy exceeded twice its own expectation, which is robust both to the third of genes with no reads in a 16-nt window and to the small number of genes that dominate pooled sums. Significance was assessed against a permutation of energy labels across genes. As a control, the same readout was applied to the opposite half of the coding sequence; because that window has a lower baseline, both readouts were additionally split at their own medians so that control and test were compared at equal power.

### Statistical analysis

rRNA hybridization profiles were obtained from. We defined strong binding sites as ones with free energy below -2.6^4,32^. We tested footprint enrichment surrounding binding sites as follows. For each strong binding site, we calculated the mean mapped footprints of size 20-45bp in the upstream / downstream 20bp. This resulted in a distribution of values, one value for each binding site. We then repeated the calculation for 100 random permutations of the strong site positions, preserving the number of sites in each region of every gene (5’-UTR, CDS, 3’-UTR). For the CDS, we considered 3 sub-regions for analysis: The first 50 base-pairs, the last 50 base-pairs, and the intermediate sub-region. We compared the average footprint occupancy (footprints / bp) in windows surrounding strong sites to the distribution of average occupancies around random sites, and computed an empirical p-value: p_emp = (|{null_avg >= real_avg}| + 1) / (n + 1) for the enrichment test, and the complementary test for the depletion test.

### Operon-junction site stratification

Strong hybridization sites within the first 8 bp of the 3′ UTR (n = 1508) were stratified by signed intergenic distance to the next downstream same-strand gene, defined as the gap between the upstream gene’s stop codon and the downstream gene’s start codon (negative when the two coding sequences overlap). The 8-bp window matches the canonical Shine–Dalgarno-to-start-codon spacing (5–13 bp^33^) expected for a downstream-gene Shine–Dalgarno positioned at this location. Sites with 0 ≤ d < 20 bp, within typical operon-internal spacing in E. coli, were classified as operon-junction sites (n = 668); sites with d ≥ 20 bp as non-junction (n = 761); and sites with d ≥ 100 bp (n = 553) provided a sharper negative control. Sites with overlapping next genes (d < 0; n = 79) were treated as a separate cohort and carried near-baseline signal, consistent with the predicted motif falling inside the next gene’s coding sequence rather than at its Shine–Dalgarno register. The Shine–Dalgarno register distribution for operon-junction sites is shown in Supplementary Figure S 3.

### Per-site enrichment test

Per-site SSU signal was quantified on the 0.5% SSU library as the mean 5′-end count of reads at the canonical Shine–Dalgarno register upstream of the start codon (read length L ∈ [30, 45) and 5′-end position P_5 ∈ [−25, −15] relative to the strong site). Because per-gene profiles concatenate the 5′ UTR, CDS, and 3′ UTR, the P_5 ∈ [−25, −15] offset from a strong 3′ UTR site (s = 0–7) reads backward across the CDS / 3′ UTR boundary into the upstream gene’s CDS-tail. This is the genomic location of the SSU footprint 5′-end when the footprint is bound to a Shine–Dalgarno motif just past the upstream stop codon. Operon-junction and non-junction signals were compared with a one-sided Mann–Whitney U test on the per-site distributions. To control for differences in gene expression between cohorts, per-site signals were normalized by gene CDS RPKM as signal / max(RPKM, 1.0), with the floor at 1 stabilizing the ratio at low expression. The resulting junction-to-non-junction enrichment was stable at 6.4–7.0× across RPKM floors of 0.1, 1.0, and 10.0 and under additive shrinkage signal / (RPKM + 1), and at 5.0–8.0× across read-length bands within the SSU-dominant range of 20–50 nt.

### Metagene profile generation

Metagene profiles were generated by averaging footprint signal across a ±30 nt window surrounding each predicted hybridization site. Coverage profiles were corrected for transcript boundaries. A read can only be placed if both its 5′ end and its 3′ end fall inside the transcript (in the 5′ UTR, inside the annotated window). For each site and displayed position, we counted the (read length, 5′-end position) combinations that could place a read covering that position. We divided the site’s observed coverage by this count, rescaled to the count available far from any boundary, with the correction capped at five-fold. A position near a transcript end is therefore compared with what could be observed there, not with what could be observed mid-gene. The correction assumes that read placements blocked by a transcript end would have been as frequent as the unblocked ones. Where fewer than about half of all placements are possible, the corrected value rests on little data and should be interpreted with caution. In the 3′ UTR the boundary is the end of the 3′ UTR window, which for some genes extends beyond the transcription terminator; positions beyond the terminator are not corrected, as the terminator is annotated only for a small fraction of genes (RegulonDB v12). The fraction of possible placements is shown beneath each profile in Figure 5.

### Correlation analysis of ribosomal footprint distributions

Spearman correlation coefficients were calculated for footprints of length 20–45 nt to assess the relationship between ribosomal footprint distributions across libraries. Correlations were computed using normalized footprint read density across transcripts, comparing SSU-, LSU-, and monosome-associated footprints under both non-cross-linked (0.0%) and 0.5% formaldehyde cross-linked conditions. Analyses were performed separately for the 5′ UTR, CDS, and 3′ UTR. Cross-linking comparisons were used primarily to evaluate footprint recovery efficiency and technical consistency, while biological interpretation focused on correlations between ribosomal populations within each transcript region.

## RESULTS

### 1. SSU-seq approach for tracking the small subunit of the ribosome in prokaryotes

SSU-seq is an advanced high-throughput sequencing technique designed to capture the positions of both assembled and partially assembled ribosomal complexes on mRNA, thereby providing a more comprehensive view of translation dynamics than conventional ribosome profiling^18^. Unlike Ribo-seq, which predominantly captures fully assembled ribosomes, SSU-seq enables the detection of small ribosomal subunits (SSU), large subunits (LSU), and monosomes, allowing direct interrogation of ribosomal subunit behavior during translation initiation, elongation, and termination. This capability provides unique insight into the stepwise engagement of ribosomal subunits with mRNA and into regulatory mechanisms acting at early stages of translation^18,23^.

In this study, we optimized the SSU-seq protocol for *E. coli*. A schematic overview of the experimental workflow is shown in Figure 1(b). Briefly, translation is arrested at mid-log growth, followed by formaldehyde-mediated cross-linking to preserve ribosomal subunit– mRNA interactions and rapid freezing of cells in liquid nitrogen. Cells are lysed using a combination of mechanical pulverization and chemical lysis, and unprotected mRNA regions are digested using micrococcal nuclease (MNase). Ribosome–mRNA complexes are subsequently separated by sucrose gradient ultracentrifugation, enabling fractionation of distinct ribosomal populations. Ribosome-protected RNA fragments are recovered by phenol extraction, converted into sequencing libraries, and subjected to high-throughput sequencing. Footprints are then mapped to the *E. coli* genome to infer ribosomal subunit positioning along transcripts^18,34^. Together, this optimized SSU-seq workflow enables high-resolution mapping of ribosomal subunits across the transcriptome, providing an experimental framework for analyzing translation initiation dynamics and rRNA–mRNA interactions in vivo. Detailed descriptions of each experimental step are provided in the Methods section.

We conducted three distinct SSU-seq experiments, generating a total of nine libraries. For each experiment, we prepared separate libraries for footprints associated with the small subunit (SSU), large subunit (LSU), and monosomes. The experiments were designed as follows and are summarized in

Figure 2a (1) A single experiment was performed without cross-linking (0% formaldehyde), mimicking the standard ribo-seq protocol (2) Two experiments were conducted using 0.5% formaldehyde. In Figure 2b–d, polysome profiles show the distribution of ribosomal complexes along a sucrose gradient, where the y-axis represents absorbance and the x-axis represents position along the tube. Lower-density particles appear at the top of the gradient (beginning of the x-axis), with density increasing along the tube; accordingly, the first peaks correspond to the SSU, followed by the LSU and then monosomes.

**Figure 2:**
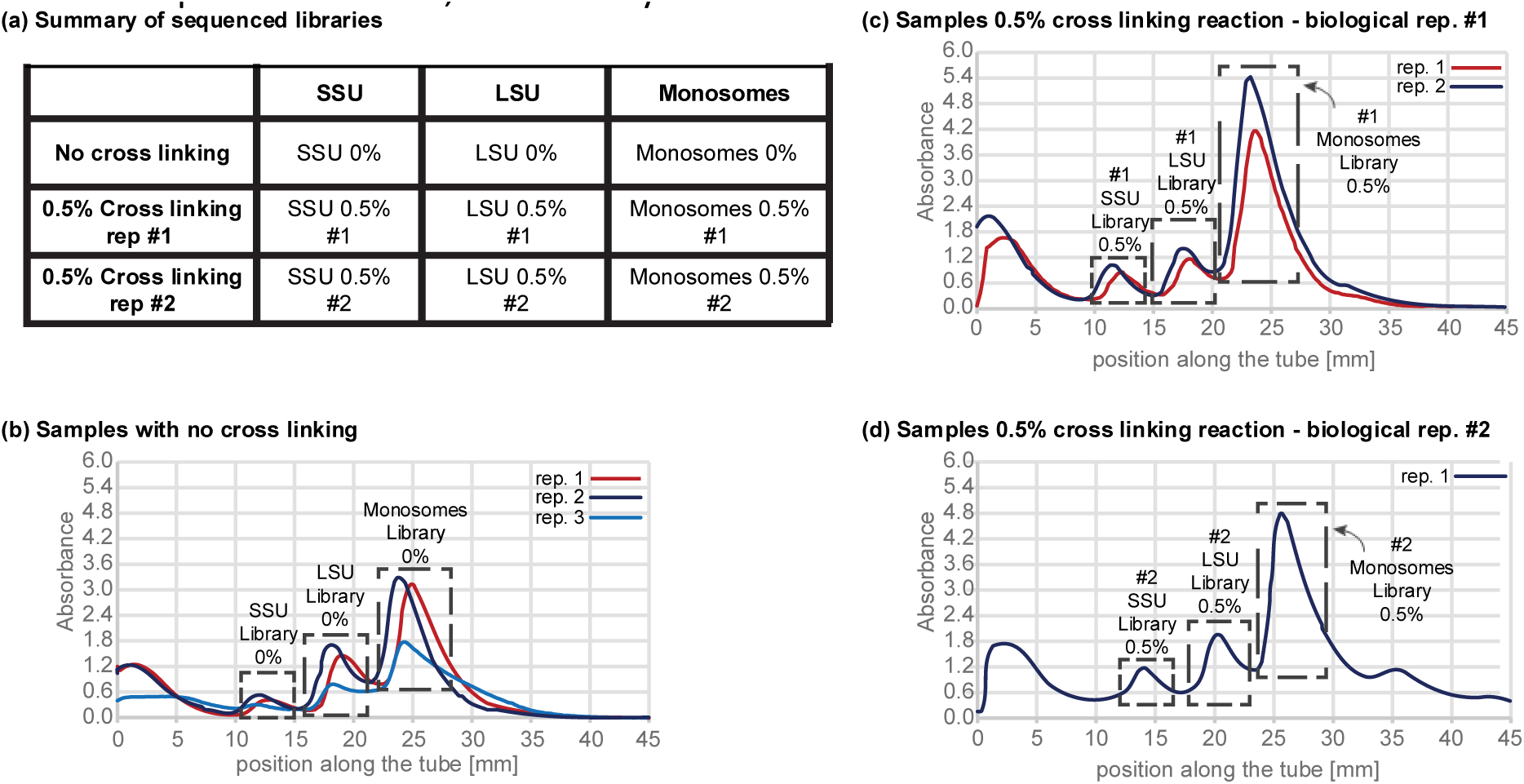
polysome profiling results of the sequenced samples (a) Table summarizing the nine libraries sequenced, including samples with 0.5% formaldehyde cross-linking and 0% formaldehyde control (b) Polysome profile of SSU-seq samples without cross-linking (0% formaldehyde) (c) Polysome profile of SSU-seq samples with 0.5% formaldehyde cross-linking, biological replication #1(d) Polysome profile of an additional biological replicate with 0.5% formaldehyde cross-linking replication #2, demonstrating reproducibility.

### 2. mRNAs with more ribo-seq reads tend to have more SSU-seq reads

We isolated footprints ranging from 15-75 bp in length. The distribution of read counts by length is shown in Figure 3 (left) for (a) SSU (b) LSU and (c) monosomes. These lengths were compared to those obtained from the ribo-seq protocol in *E. coli*. The Spearman’s correlation between our data and the ribo-seq data^24^ (Figure 3,right) indicates a relatively high correlation (>0.6) for SSU footprints of 15-50nt and monosomes footprints of 20-50nt. Differences in biological protocols may account for reduced correlation in read numbers and lengths.

**Figure 3:**
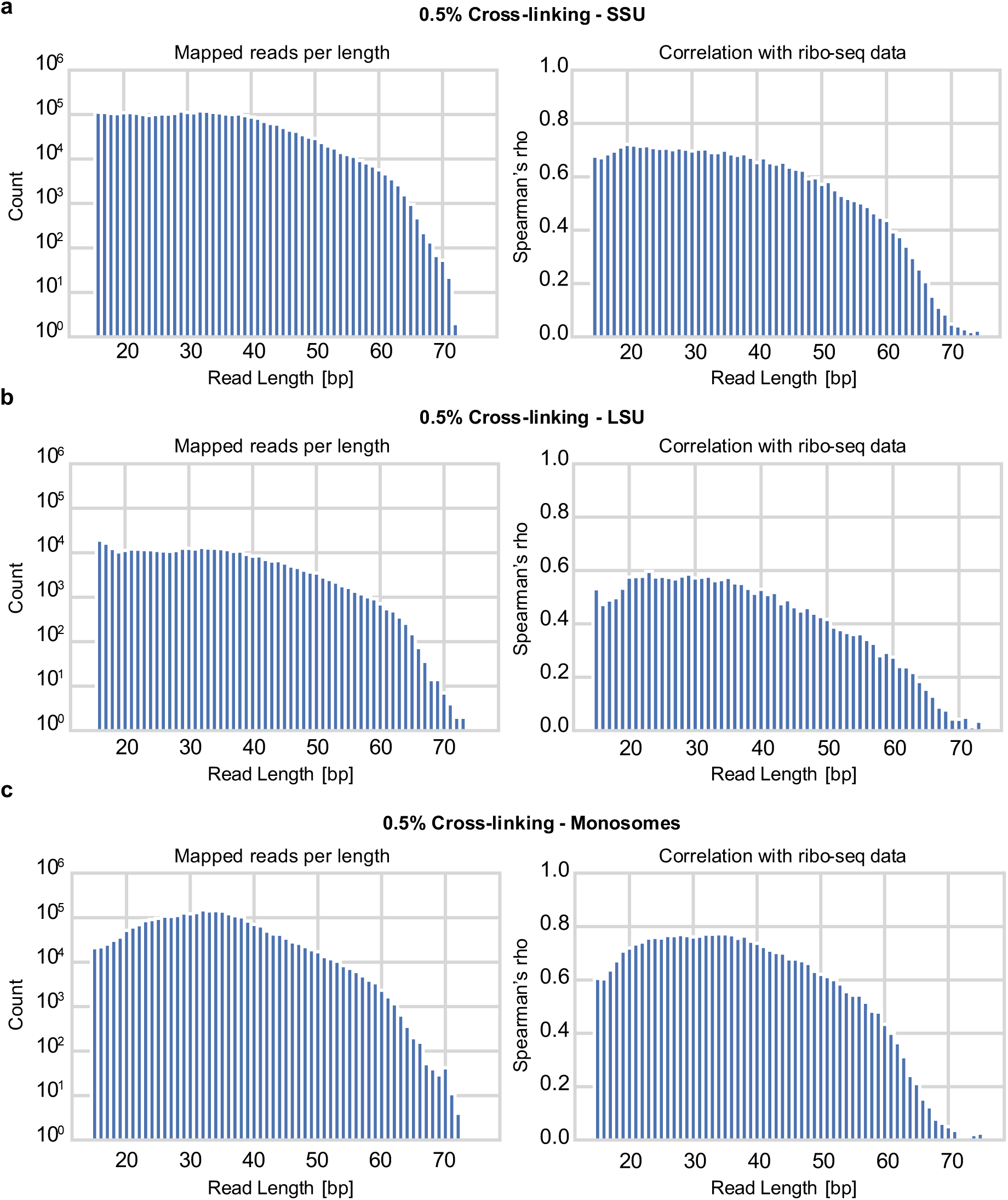
Analysis of ribosome-protected mRNA fragment lengths in SSU-seq compared to Ribo-seq. On the left distribution of read counts across different footprint lengths (15-75 bp) obtained from SSU-seq. On the right Spearman’s correlation between SSU-seq and Ribo-seq data plotted against read length. The x-axis represents footprint length in nucleotides, and the y-axis shows the correlation coefficient while (a) represents footprints of SSU (b) footprints of LSU and (c) footprints of the monosomes

### 2.1. Cross-Linking Enables Recognition and Isolation of Ribosomal Subunits-Protected mRNA Footprints near rRNA-mRNA hybridization site

The following results are based on a dataset derived from samples treated with 0.5% formaldehyde. Set 1 corresponds to sites with strong rRNA-mRNA hybridization and Set 2 to sites with intermediate rRNA-mRNA hybridization (Figure 4). Figure 4a illustrates the positional framework used for footprint mapping relative to the first nucleotide of the predicted hybridization site. Additional dataset of untreated samples is presented in the supplementary section, Figure S 1 and S 4.

**Figure 4:**
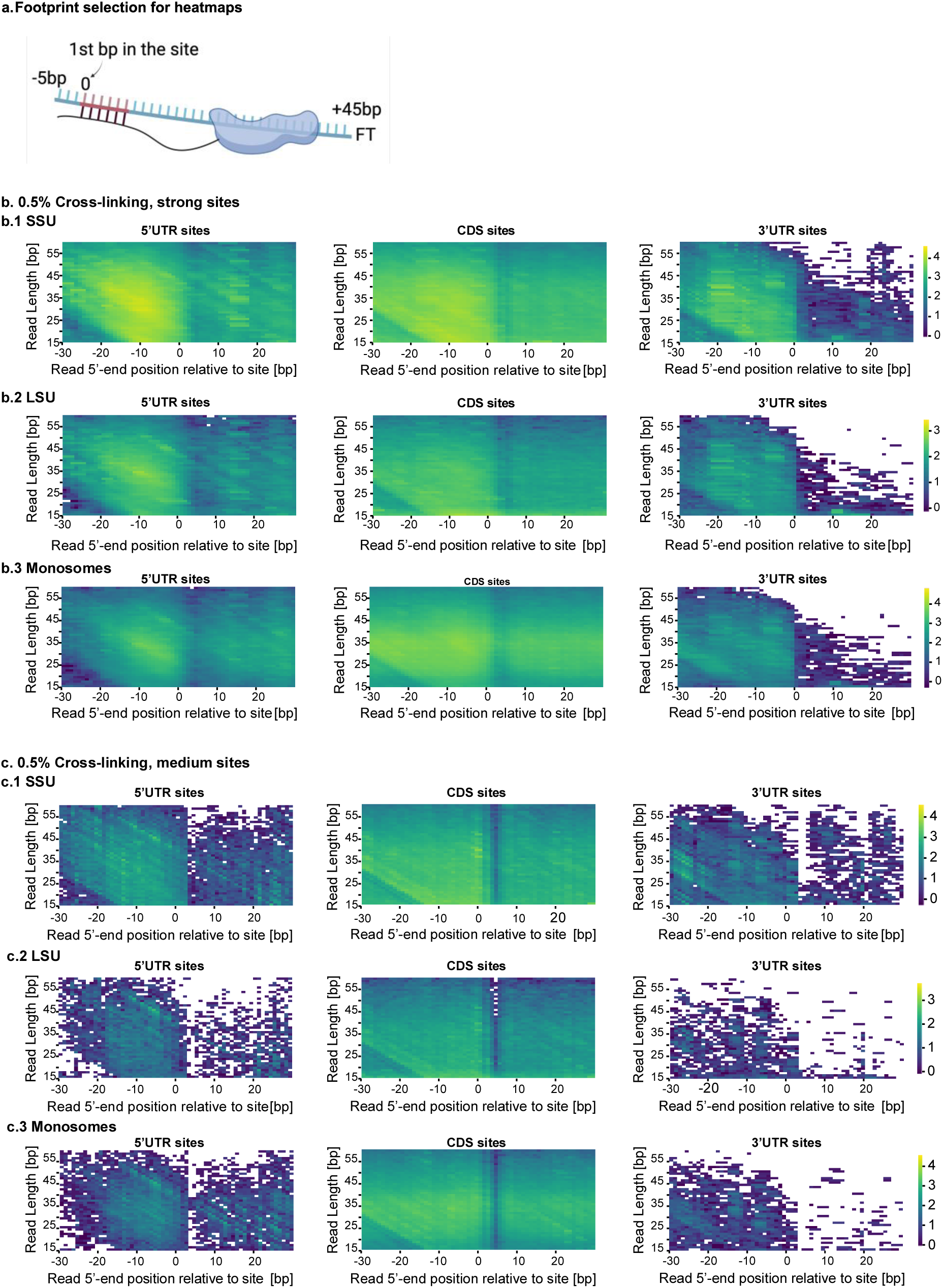
Heatmap analysis of ribosomal subunit and monosome binding patterns across mRNA transcripts obtained by SSU-seq using 0.5% formaldehyde. (a) Schematic representation of the footprint (FP) populations considered for heatmap construction. An illustrative example of a 50 bp footprint spanning from 5 bp upstream to 45 bp downstream of a strong/intermediate site is shown. SSU is presented as an example, but the same principle was used for LSU and monosome footprints in the heatmap analysis. (b) Heatmaps corresponding to strong rRNA–mRNA hybridization sites. (c) Heatmaps corresponding to intermediate rRNA–mRNA hybridization sites. Heatmaps illustrate the distribution of ribosome-protected mRNA fragments across transcript regions: the 5′ untranslated region (5′ UTR, left), coding sequence (CDS, middle), and 3′ untranslated region (3′ UTR, right). Each row represents a different ribosomal species: (1) small subunit (SSU), (2) large subunit (LSU), and (3) monosomes. Color intensity (yellow) reflects the relative frequency of fragment mapping at each position. Additional datasets and biological replicates are presented in Supplementary Figure S 4. In the 3′ UTR panels, the reduced signal at long read lengths and downstream displayed positions reflects genuinely lower read coverage downstream of 3′ UTR hybridization sites: small subunit occupancy declines with distance past the stop codon, and, because the 3′ UTR window is a fixed length and does not mark the transcript end, no reads can map beyond the transcription terminator where it falls inside the window. The 3′ UTR also carries fewer predicted hybridization sites overall than the other regions. The 5′ UTR panels show a milder, length-independent reduction at upstream displayed positions: for sites positioned within the first ∼30 bp of the 5′ UTR, very negative window positions would map past the transcript’s 5′ boundary and so cannot contribute at those cells.

In both sets, a high number of footprints were recovered for all ribosomal components, indicating efficient cross-linking of ribosomes and ribosomal subunits to the mRNA near rRNA-mRNA hybridization site. The SSU-associated signal is especially enriched in the 5′ UTR, approximately 8–10 nucleotides upstream of the start codon, consistent with the ribosome binding site. Additionally, strong signals are observed within the CDS at sites of robust rRNA-mRNA hybridization, suggesting SSU engagement with these coding regions as well.

The LSU binding pattern in both the strong-site dataset (Figure 4 – b2) and the intermediate-site dataset (Figure 4 – c2) shows weaker signals compared to the SSU in both the 5′ UTR and CDS. The signal in the 5′ UTR may partly result from dissociated monosomes initially bound at the RBS. Signal in the 3′ UTR is minimal, likely due to a lower number of mapped reads in that region. Monosome binding (Figure 4 – b3, c3) displays a weak but distinct signal near the RBS in the 5′ UTR, and a more pronounced signal across the CDS. Together, the results from both intermediate and strong rRNA-mRNA hybridization sites highlight a consistent pattern of SSU interaction with the 5′ UTR and CDS, and demonstrate the importance of cross-linking for stabilizing ribosome-mRNA associations.

### 2.2. Evidence for SSU Association with CDS and 3′ UTR Regions Driven by rRNA-mRNA Base Pairing

Mean footprint coverage was calculated within a ±30 bp window centred on predicted rRNA–mRNA hybridization sites (Figure 5a) of intermediate and strong strength (Figure 5b) across the 5′ UTR, CDS, and 3′ UTR. Panels 1–3 show SSU-, LSU-, and monosome-derived footprints, respectively. Data correspond to the 0.5% formaldehyde cross-linked sample. The x-axis denotes the position relative to the first nucleotide of the predicted hybridization site (position 0), and the y-axis represents normalized footprint coverage. Additional samples generated without cross-linking (0% formaldehyde; Figure S6) show qualitatively similar but weaker enrichment patterns, supporting the interpretation that cross-linking enhances recovery of transient SSU–mRNA interactions without altering overall ribosome distribution.

**Figure 5:**
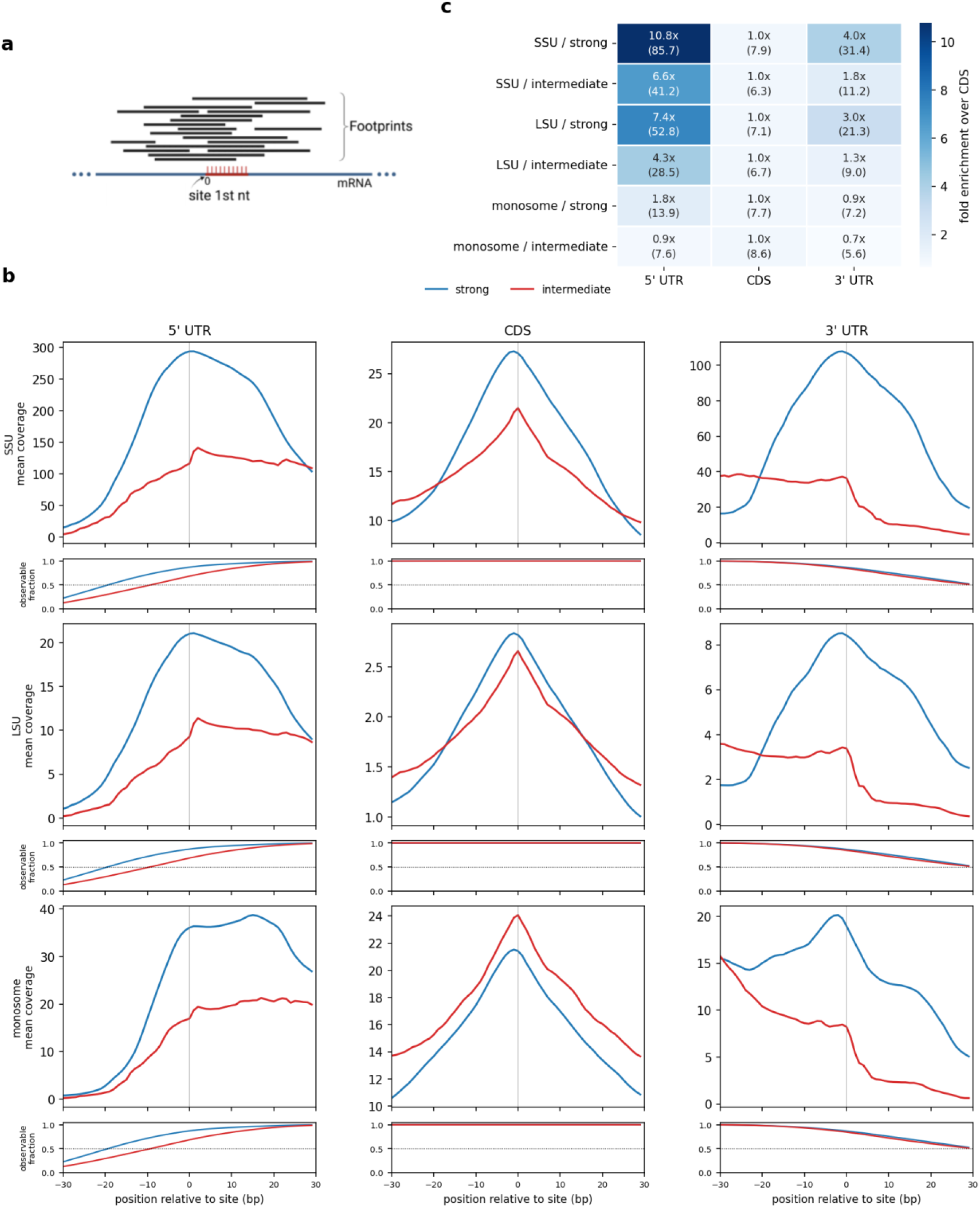
Mean ribosome-protected footprint coverage around predicted rRNA–mRNA hybridization sites across mRNA regions. Data are shown for the 0.5% formaldehyde cross-linked sample. (a) Schematic illustration of footprint alignment relative to predicted rRNA–mRNA hybridization sites; position 0 corresponds to the first nucleotide of each site. (b) Mean normalized footprint coverage within a ±30 bp window centered on sites with strong (blue) / intermediate (red) predicted hybridization strength (n = 14,477 / 119,503 / 7,096 strong and 1,618 / 46,315 / 1,641 intermediate, for 5′ UTR / CDS / 3′ UTR). Coverage at each displayed position is normalized by the number of (read length, 5′-end position) combinations that could place a read there (see Methods), so that profiles are comparable near transcript boundaries. The strip beneath each profile shows the fraction of combinations possible at that position. Rows correspond to SSU-, LSU-, and monosome-associated footprints (top to bottom). The x-axis represents the position relative to the hybridization site, and the y-axis indicates normalized footprint coverage. (c) Heatmap summarizing peak footprint enrichment across mRNA regions and hybridization strengths. Values are shown as fold enrichment relative to the CDS peak, with RPM-normalized peak coverage (per million mapped reads) indicated in parentheses. Fold enrichment values are library-size invariant. Additional datasets and biological replicates are presented in Supplementary Figure S 6.

SSU-, LSU-, and monosome-derived footprints exhibit distinct regional distributions. SSU footprints are detected across all transcript regions, with pronounced enrichment near specific hybridization sites. Monosome footprints show strong and continuous coverage along the CDS, consistent with elongating ribosomes and largely independent of hybridization strength. In contrast, LSU-associated footprints display weaker and less structured distributions across regions.

Each hybridization site was defined at single-nucleotide resolution. The binding matrix described by Bahiri et al. (2021)^5^ assigns a predicted rRNA–mRNA hybridization energy to every nucleotide along the transcript. Although complementarity is computed over a fixed window (corresponding to the site length selected in the original model), the output is resolved per nucleotide position. Positions were classified according to predicted hybridization strength using predefined thresholds; strong sites: ΔG < –2.6, intermediate sites: –2.6 ≤ ΔG < –1.7 (see more details in the methods section).

Metagene profiles revealed distinct positional enrichment patterns across transcript regions and between strong and intermediate rRNA–mRNA interaction sites.

In the 5′ UTR, both site types display a leftward ramp toward the start codon, peaking around positions 0–5 bp in the SSU and LSU profiles; monosome profiles instead form a broad plateau that peaks 15–17 bp downstream of the site. However, profiles centered on intermediate sites are broader and flatter. Small subunit footprints are positioned by the canonical Shine–Dalgarno sequence rather than by each predicted hybridization site, and intermediate-strength sites span a wider range of distances to the start codon than strong sites (median 23 nt versus 13 nt). A profile centered on a site further upstream therefore shows the same footprint displaced downstream by the distance from that site to the Shine–Dalgarno sequence, so the width of the profile reflects the spread of this distance across the cohort. Consistent with this, the footprint signal at the displaced position exceeded the signal at the site itself by 3.3-fold to 14.9-fold for intermediate sites 30 to 40 nt from the start codon, and by 3.1-fold to 22.8-fold for strong sites at the same distances (95% bootstrap intervals excluding 1 in all three subunit fractions).

In the 3′ UTR, strong sites show a clear central peak, resembling a mirrored pattern of the 5′ UTR. In contrast, intermediate sites lack a defined peak and instead display a gradual decay from the CDS boundary toward the transcript end.

Within coding sequences, both strong and intermediate sites show only modest enrichment near positions –5 to 0. For small subunit footprints, peak coverage was nonetheless higher at strong than at intermeidate sites (8 7.9 versus 6.3 RPM; Figure 5c

), a smaller difference than in the 5′ UTR but in the same direction. Monosome footprints showed no such difference (7.7 versus 8.6 RPM), consistent with their coding-sequence signal reflecting elongating ribosomes rather than hybridization-dependent recruitment. Enrichment in the 5′ UTR is substantially higher in absolute terms, indicating that the regions differ in the overall level of small subunit engagement rather than in whether predicted hybridization strength is reflected at all.

To enable cross-region comparison, enrichment was quantified relative to the CDS peak. SSU footprints show the strongest enrichment in the 5′ UTR, reaching ∼11-fold over CDS at strong sites and ∼7-fold at intermediate sites, consistent with strength-dependent recruitment upstream of the start codon. LSU footprints display a similar but less pronounced trend.

In contrast, monosome footprints show minimal 5’ UTR enrichment (1.8-fold at strong sites; ∼1-fold at intermediate sites), consistent with their primary association with elongating ribosomes. Across CDS regions, RPM-normalized peak coverage is comparable among SSU (7.9), LSU (7.1), and monosome (7.7) footprints, supporting the interpretation that CDS signal largely reflects translating ribosomes rather than hybridization-dependent recruitment. 3’ UTR enrichment is detectable but weaker than in 5’ UTR, with SSU enrichment reaching 4.0-fold at strong sites and 1.8-fold at intermediate sites.

To investigate the origin of the strong-site SSU enrichment observed in the 3′ UTR, we analyzed strong rRNA–mRNA hybridization sites located within the first 8 bp of the 3′ UTR according to their signed intergenic distance to the next downstream same-strand gene (see Methods). Sites positioned at operon junctions (0 ≤ d < 20 bp) displayed substantially higher SSU occupancy at the Shine–Dalgarno register than non-junction sites (d ≥ 20 bp), with an approximately 20-fold higher raw signal. This enrichment remained significant after normalization by gene CDS expression levels, retaining an approximately 7-fold higher signal.

SSU occupancy decreased progressively with increasing intergenic distance and was strongly concentrated within the 0–20 bp range corresponding to canonical Shine–Dalgarno spacing. In contrast, distant non-junction sites and strong sites located within CDS regions displayed minimal signal at the Shine–Dalgarno register. Together, these results indicate that the strong-site SSU enrichment observed in the 3′ UTR is primarily concentrated at canonical-spacing operon-junction positions, where the predicted hybridization site likely functions as the Shine–Dalgarno sequence of the downstream gene rather than as a feature of the upstream gene’s 3′ UTR. These findings are consistent with previous reports describing widespread SD-like motifs within bacterial 3′ UTRs, while further demonstrating that strong SSU occupancy is preferentially associated with operon-junction contexts. Additional statistical analyses, normalization controls, and negative-control comparisons are presented in Supplementary Figure S 5.

To control for differences in gene composition across regions, we repeated the analysis on a gene-matched subset in which each gene contributed an equal number of hybridization sites to every region. Enrichment of SSU footprints in the 5′ UTR at strong sites persisted under this control, at reduced magnitude, indicating that the positional enrichment is not solely a consequence of which genes contribute sites to each region.

To test whether the strength dependence reported above reflects predicted complementarity rather than proximity to the start codon, small subunit occupancy was measured at a fixed position relative to the start codon and compared against per-gene predicted hybridization energy in the canonical Shine–Dalgarno register. Occupancy scaled monotonically with predicted energy across all five quintiles and all three footprint fractions, from 70.1% of genes above twice their own expectation in the strongest quintile to 27.1% in the weakest for the small subunit (Spearman ρ = −0.29, p = 1.3 × 10⁻³⁰; z = −11.0 against permuted energy labels; large subunit ρ = −0.33, monosome ρ = −0.29; Figure S 7a). Because every gene contributes at the same position, this relationship cannot arise from differences in distance to the start codon. The same measurement applied to the opposite half of the coding sequence was flat in all three fractions when both readouts were compared at equal power (Spearman ρ = +0.03, +0.05 and +0.01, all non-significant; small subunit shown in Figure S 7b). The relationship persisted within every tercile of the readout window’s GC content, indicating that it does not arise from sequence composition.

While the site-centered analysis above highlights strength-dependent enrichment at predicted hybridization sites, we next examined global footprint distributions independent of site classification to assess overall regional behavior and reproducibility across experimental conditions (Figure 6). The global density analysis revealed that SSU-associated footprints display region-dependent and heterogeneous length distributions, whereas monosome-associated footprints exhibit a conserved and uniform profile across transcript regions. We therefore examined whether these differences are reflected in the reproducibility of footprint distributions across experimental conditions using correlation analysis.

**Figure 6:**
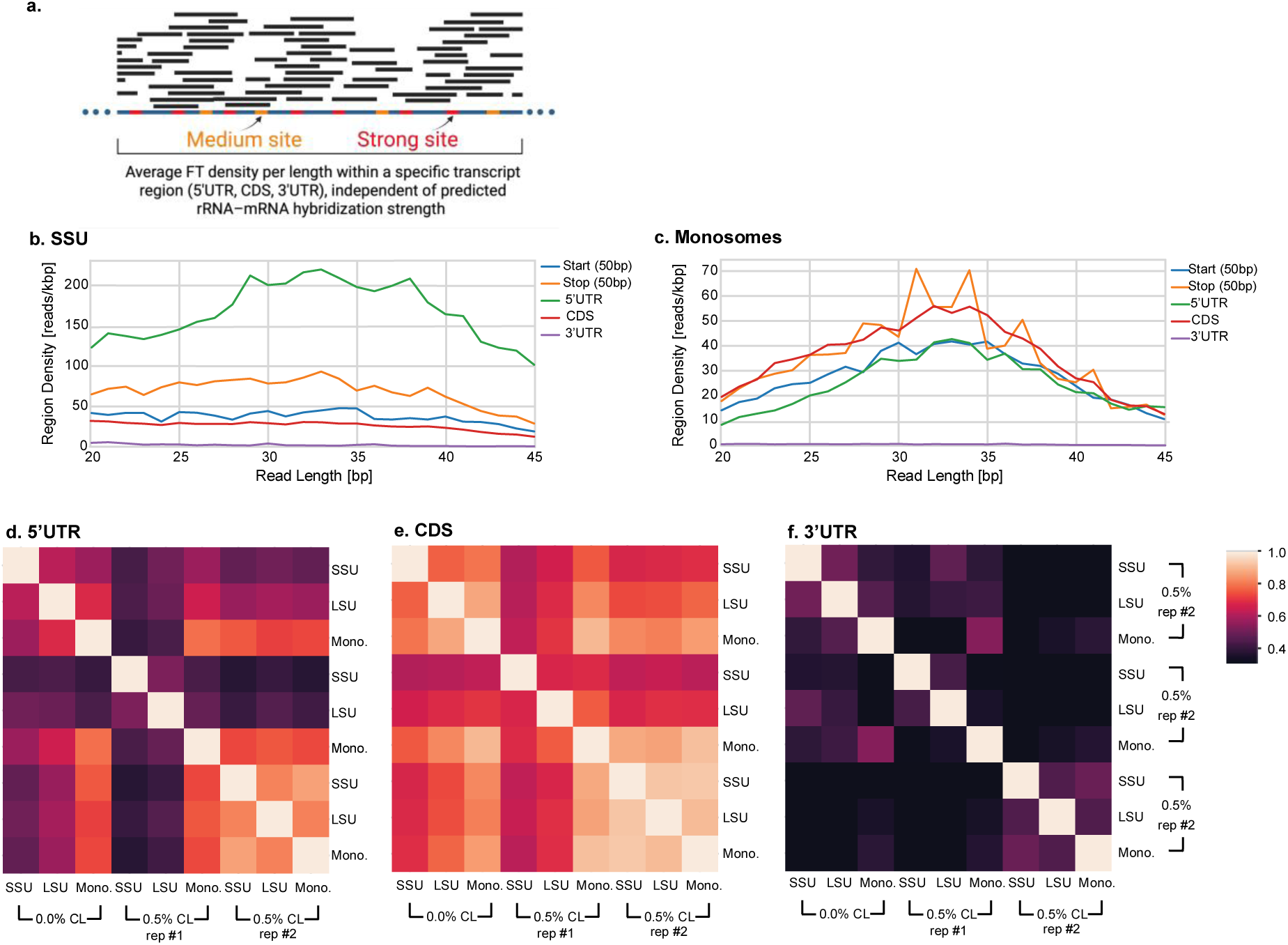
Global density and correlation analysis of SSU- and monosome-associated ribosomal footprints. (a) Schematic illustration of footprint density calculation. Ribosome footprints of varying lengths mapped to mRNA transcripts were used to calculate average footprint density around predicted strong and intermediate rRNA–mRNA hybridization sites. (b–c) Global footprint density distributions as a function of read length and transcript region, calculated without conditioning on predicted rRNA–mRNA hybridization sites. (b) SSU-associated footprints and (c) monosome-associated footprints. Color intensity represents normalized footprint density within each region. (d– f) Spearman correlation analysis of ribosomal footprint libraries (20–45 nt) generated without cross-linking (0.0%) and with 0.5% formaldehyde cross-linking. Correlations were computed based on normalized footprint read densities across transcripts, and are shown separately for the 5′ UTR (d), CDS (e), and 3′ UTR (f), and include comparisons between SSU-, LSU-, and monosome-associated libraries. Heatmap color intensity indicates the strength of correlation, with higher values reflecting greater similarity between footprint density distributions.

Spearman correlation analysis was performed for footprints of length 20–45 nt to examine the relationship between SSU-, LSU-, and monosome-associated footprint distributions across transcript regions (5′ UTR, CDS, and 3′ UTR; Figure 6d–f). Correlations were calculated based on normalized footprint read density along transcripts. Overall, positive correlations between SSU and monosome footprints were observed across transcript regions, with the strongest associations in the 5′ UTR and weaker correlations in the CDS and 3′ UTR.

In the 5′ UTR, SSU footprints show strong positive correlations with monosome-associated footprints, indicating that regions enriched in initiating ribosomal small subunits are also enriched in assembled ribosomes. This suggests coordinated ribosome recruitment and initiation, where efficient SSU binding is associated with increased monosome occupancy. Within the CDS, correlations between SSU and monosome footprints remain positive, although generally weaker than in the 5′ UTR. This may reflect translational regions with increased ribosome occupancy or local ribosome traffic, where both scanning or transiently bound small subunits and elongating ribosomes accumulate. Such patterns may indicate ribosome pausing or traffic jams associated with translational regulation that involve both ribosomes and SSU (i.e. cases where ribosomes that block SSUs and vice versa).

In the 3′ UTR, correlations are weaker overall, consistent with reduced ribosome occupancy in this region following translation termination and ribosome dissociation. Nevertheless, detectable associations between SSU and monosome footprints suggest that ribosomal subunits may remain transiently associated with these regions.

## DISCUSSION

This study presents an optimized SSU-seq protocol for *E. coli* that integrates formaldehyde cross-linking to stabilize ribosome–mRNA interactions, enabling the isolation and analysis of footprints derived from small ribosomal subunits, large subunits, and monosomes. By integrating regional stratification with hybridization-strength classification, we establish a quantitative, genome-wide framework for resolving small subunit positioning and rRNA–mRNA interaction landscapes beyond the scope of conventional Ribo-seq.

### Validation of SSU-seq and Cross-Linking Dependence

Analysis of footprint lengths demonstrated that SSU and monosome fragments fall within expected size ranges and show strong Spearman correlation with previously published Ribo-seq datasets. SSU footprints (15–50 nt) and monosome (20–50 nt) footprints were particularly well correlated, supporting the robustness of the protocol in capturing biologically relevant ribosome–mRNA complexes. The agreement between monosome occupancy and prior datasets provides independent validation of the SSU-seq approach. Because assembled ribosomes mediate productive translation, regions enriched in monosome footprints are expected to coincide with increased small subunit occupancy. The observed consistency between SSU and monosome signals is therefore biologically consistent and suggests that SSU-seq captures authentic translational states rather than experimental artifacts.

In contrast, LSU footprints exhibited weaker correlation and broader length distributions, suggesting that LSU–mRNA interactions are more transient or less efficiently stabilized under the conditions used. One possible innovative explanation is that the large ribosomal subunit can engage mRNA independently of fully assembled ribosomes. Such interactions may be shorter-lived, or structurally heterogeneous, resulting in broader footprint distributions and reduced reproducibility. Alternatively, LSU-associated signals may reflect transient assembly intermediates or post-termination states that are inherently more variable.

Consistent with this interpretation, subunit-associated footprints were markedly reduced in the absence of cross-linking, emphasizing the dynamic nature of subunit–mRNA interactions and the necessity of cross-linking for capturing early or partial ribosomal engagement states in vivo.

### Strength-Dependent SSU Enrichment and Regional Behaviour

Site-centered analysis revealed clear strength-dependent enrichment of SSU footprints, particularly within the 5′UTR. Strong predicted hybridization sites exhibited sharp enrichment peaks, whereas profiles centered on intermediate sites were broader, reflecting their wider distribution of distances to the start codon. Within the coding sequence the same intermediate sites show enrichment at the site itself, where no dominant upstream Shine–Dalgarno sequence anchors the footprint elsewhere: the two regions differ in what sets the footprint’s position, not in the sites themselves. Because footprint occupancy reports residence rather than productive initiation, higher occupancy at stronger predicted sites is expected under this model: an interaction that retains the subunit longer at a given position leaves a larger footprint there, whether or not that position is productive. In contrast, Monosome footprints were largely independent of hybridization strength and were enriched predominantly within the CDS, consistent with their role in elongation.

SSU footprints were detected across all transcript regions, including the 5′ UTR, CDS, and 3′UTR. Quantitative comparison using CDS-normalized fold enrichment demonstrated that 5’UTR enrichment is substantially higher in absolute terms than CDS enrichment, while a dependence on predicted hybridization strength was present in both regions. These observations indicate that enrichment in untranslated regions reflects site-specific interactions rather than differences in transcript abundance.

### A Two-Component Model of SSU–mRNA Interaction

The observed enrichment profiles can be interpreted within a two-component framework ccomprising a regional background signal and a site-specific recruitment component. The background reflects general ribosome behavior within each transcript region, such as SSU movement in the 5′UTR or progressive dissociation in the 3′UTR, whereas the site-specific component arises from rRNA–mRNA complementarity that stabilizes the small subunit at discrete positions. In the 5′ UTR, a broad background signal is superimposed with a dominant peak at strong Shine–Dalgarno-like sites; profiles centered on intermediate sites appear flatter because those sites lie at a wider range of distances from that peak. Consistent with this, small subunit occupancy at the Shine–Dalgarno register scales continuously with predicted hybridization energy across genes, rather than only between the discrete strength classes used here. The presence of this extended upstream signal suggests that, in addition to canonical Shine–Dalgarno recognition, prokaryotic initiation may involve a degree of local scanning or guided exploration. Although bacterial initiation is classically described as a direct recruitment process, these results are consistent with a model in which small subunits transiently sample nearby regions prior to start codon selection, analogous in principle to eukaryotic scanning but likely constrained to shorter distances.

In the 3′ UTR, the dissociation gradient dominates at intermediate sites, whereas strong sites generate a detectable peak above this background. Stratification by intergenic distance to the next downstream same-strand gene indicates that this strong-site peak is concentrated at operon-junction positions (approximately 7-fold over non-junction strong sites after expression normalization; p < 1e−47), where the predicted hybridization site corresponds to the Shine–Dalgarno sequence of the downstream gene. The strong-site 3′ UTR signal therefore largely reflects SSU recruitment at the downstream gene’s Shine–Dalgarno sequence rather than persistent association with the upstream gene’s terminator region. The intermediate-site 3′ UTR signal, by contrast, follows a broader dissociation-gradient pattern consistent with post-termination SSU persistence beyond the stop codon. The much smaller difference in CDS baseline signal between strong and intermediate site sets further supports the interpretation that these differences arise from hybridization-dependent stabilization rather than transcript abundance.

### Implications for SSU Persistence and Facilitated Search

Together, these findings support a model in which the small ribosomal subunit remains associated with mRNA beyond canonical initiation regions. Rather than binding exclusively at the Shine–Dalgarno sequence and dissociating immediately after initiation, the SSU appears capable of engaging transcripts across coding and downstream regions in a hybridization-dependent manner.

This behaviour is consistent with models in which small subunits “diffuse” in proximity to mRNA and are guided by base-pairing potential. Strong hybridization may transiently trap the subunit, whereas intermediate interactions may enable repeated binding and release events that maintain proximity without trapping the it in nonproductive states. Such interactions may facilitate start-codon recognition through a guided search mechanism and contribute to efficient recycling or re-initiation following termination.

The persistence of SSU footprints in the CDS, together with the broader 3′ UTR dissociation gradient at intermediate sites, challenges the classical view that the SSU dissociates immediately after initiation or termination, and instead suggests that weak yet stabilizing rRNA-mRNA interactions contribute to shaping translation dynamics at multiple stages of the translation cycle. The strong-site 3′ UTR peak does not contribute to this evidence; its reattribution to operon-junction Shine-Dalgarno sequences (see Two-Component Model) places it on the downstream gene’s pre-initiation register rather than the upstream gene’s post-termination 3′ UTR.

### Broader Implications and Future Directions

The SSU-seq framework establishes a dedicated experimental platform for dissecting early and late stages of translation that remain inaccessible to conventional Ribo-seq. Future applications may include analysis of initiation factor mutants, recycling-defective strains, stress conditions, or diverse bacterial species. Combining SSU-seq with perturbation-based or time-resolved approaches will enable ffurther dissection of subunit dynamics under varying physiological conditions. By enabling transcriptome-wide mapping of small subunit engagement, this approach provides new insights into the spatial, kinetic, and regulatory organization of bacterial translation.

## ACKNOWLEDGEMENTS

We would like to thank Dr. Daniel Dovrat for his invaluable advice and assistance. Figures and schematic illustrations were created, in part, with BioRender (https://BioRender.com).

## AUTHOR CONTRIBUTIONS

Larissa Fine: Conceptualization, Methodology, Validation, Visualization, Writing—original draft, Visualization. Alon Diament: Formal analysis, Visualization. Rachel Cohen-Kupiec: Conceptualization, Project administration, Methodology, Formal analysis. Tamir Tuller: Supervision, Conceptualization, Methodology. Writing—original draft, Writing—original draft, Visualization. All authors – review and approve the paper.

## DATA AVAILABILITY

The data underlying this article are available in the NCBI Gene Expression Omnibus (GEO), and can be accessed with accession number GSE334537 at https://www.ncbi.nlm.nih.gov/geo/query/acc.cgi?acc=GSE334537. We deposited the raw single-end reads and the processed footprint counts for all nine SSU-seq libraries: the small subunit, large subunit and monosome fractions from each of three experiments, one performed without cross-linking (0% formaldehyde) and two with 0.5% formaldehyde as biological replicates. Each processed file gives 5’-end counts per gene, transcript region (5’ UTR, CDS, 3’ UTR), position and read length, using the Escherichia coli NC_000913.3 annotation. The record is private until publication and will be released on acceptance;

We also used two published datasets. The rRNA-mRNA hybridization profiles that define the predicted binding sites are from Bahiri Elitzur et al.. The E. coli ribosome profiling data used for the read-length correlation in Figure 3 are available in GEO under accession GSE72899.

