## Supplementary information for "Monitoring the translation initiation process in prokaryotes via large-scale examination using optimized SSU-seq technique"

#### Optimization of growth and cross-linking conditions for SSU-seq

Supplementary Figure S 1 summarizes optimization experiments described in the main text, including comparisons of growth media (LB vs. MOPS), cross-linking pH conditions, formaldehyde concentrations, and MNase digestion parameters, evaluated using RNA extraction and polysome profiling.

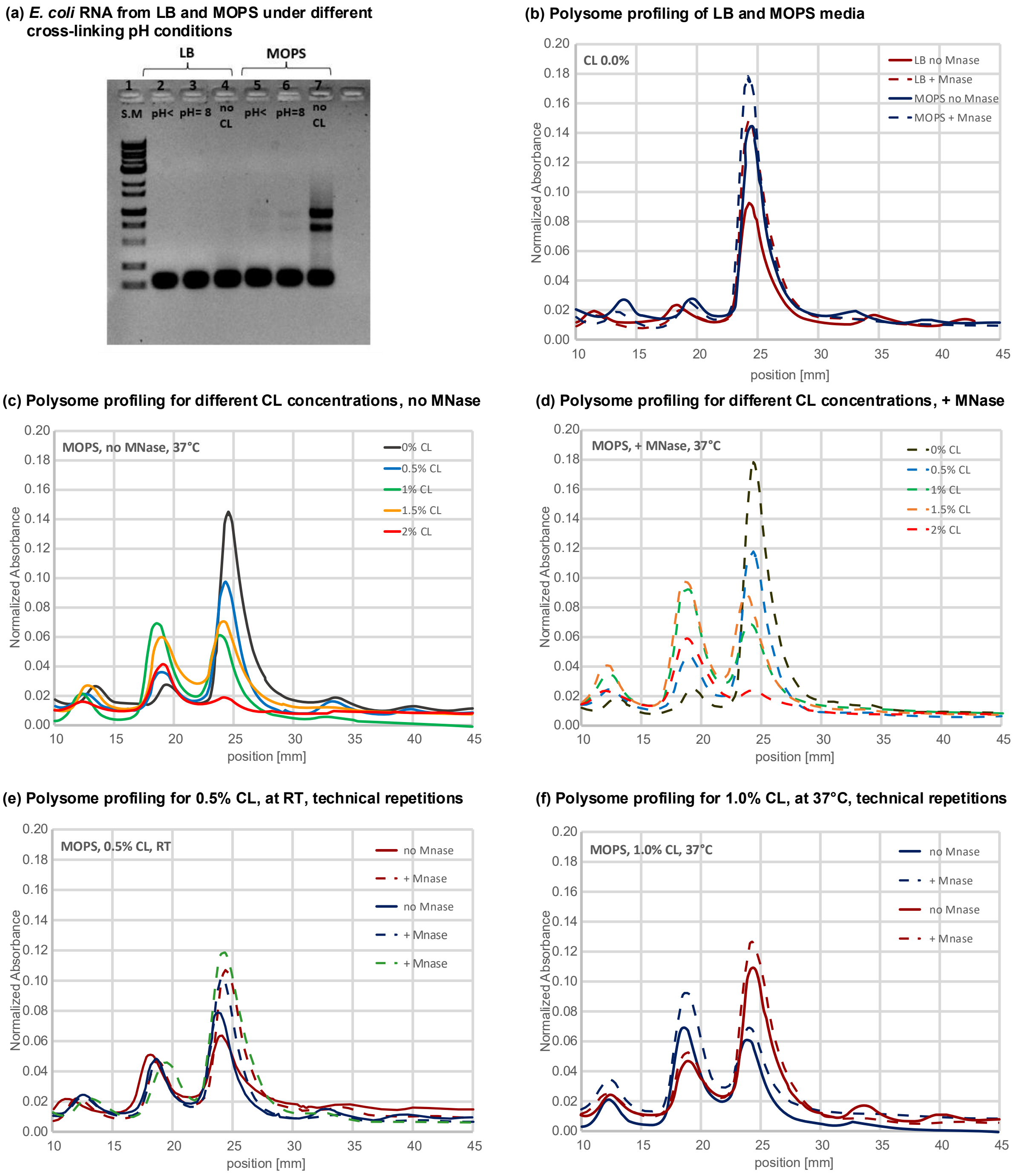


*Figure S 1: **Optimization of different parameters** (a) RNA extracted from E. coli cultures grown in LB or MOPS media, subjected to three cross-linking conditions: low pH (<5), pH 8, and no cross-linking. (b) Polysome profiling of samples grown in LB (red) or MOPS (blue) media, with (dashed) and without*

*(solid) MNase digestion. (c) Polysome profiling at different CL concentrations in MOPS at 37°C, without MNase. (d) Polysome profiling at different CL concentrations in MOPS at 37°C, with MNase. (e) Polysome profiling of samples cross-linked with 0.5% CL at room temperature, with (dashed) and without (solid) MNase. (f) Polysome profiling of samples cross-linked with 0.5% CL at 37°C, with (dashed) and without (solid) MNase.*

**Ribosome-protected fragment purification and library quality assessment**

Supplementary Figure S 2  shows the purification and quality assessment of ribosome-protected fragments following cross-link reversal and gel extraction. RNA fragments corresponding to 15–70 nt were excised from denaturing PAGE gels and analyzed using the Agilent 2100 Bioanalyzer to confirm fragment size distribution and library quality.

**(a)** ***Ribosome protected fragments purificatoin***

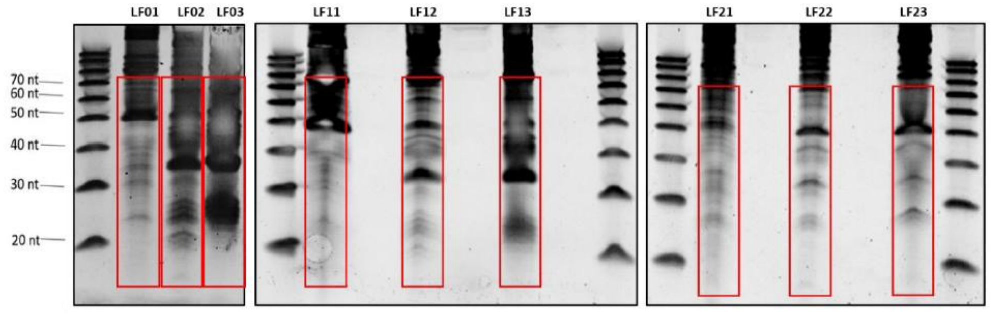


**(b)** ***Agilent Bioanalyzer profiles of libraries***

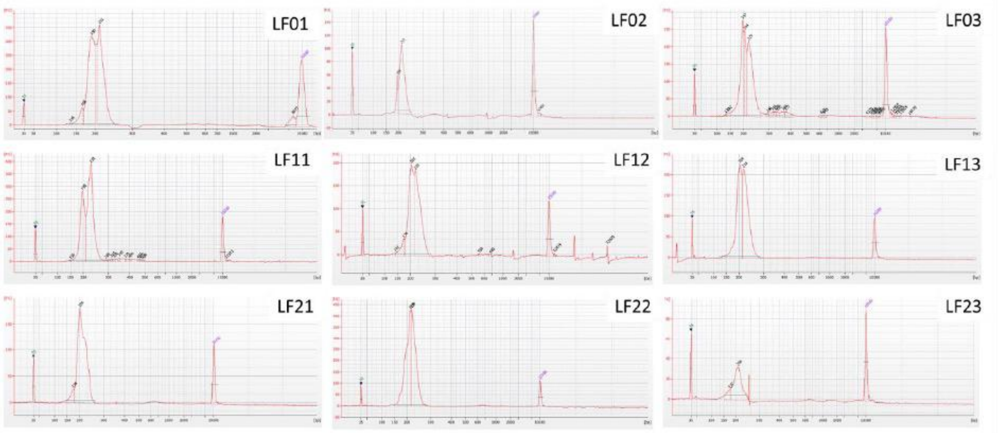


*Figure S 2:* ***Ribosome-protected fragment purification and quality assessment.***
*(a) Denaturing 15% TBE-urea gel showing RNA fragments following cross-link reversal and phenol-chloroform extraction. Red boxes indicate gel regions (15–70 nt) excised for footprint recovery. Lane labels correspond to fractions collected from polysome profiling.*
*(b) Agilent 2100 Bioanalyzer profiles of recovered RNA libraries (LF01–LF23), confirming fragment size distribution and quality of ribosome-protected fragments used for library preparation.*

**Operon-junction site stratification.** Figure S 2 further examines the positional organization of strong 3′ UTR hybridization sites classified as operon-junction sites. The analysis shows that predicted motifs are preferentially positioned at distances consistent with canonical Shine–Dalgarno spacing relative to the downstream start codon, supporting their interpretation as downstream-gene translation initiation signals.

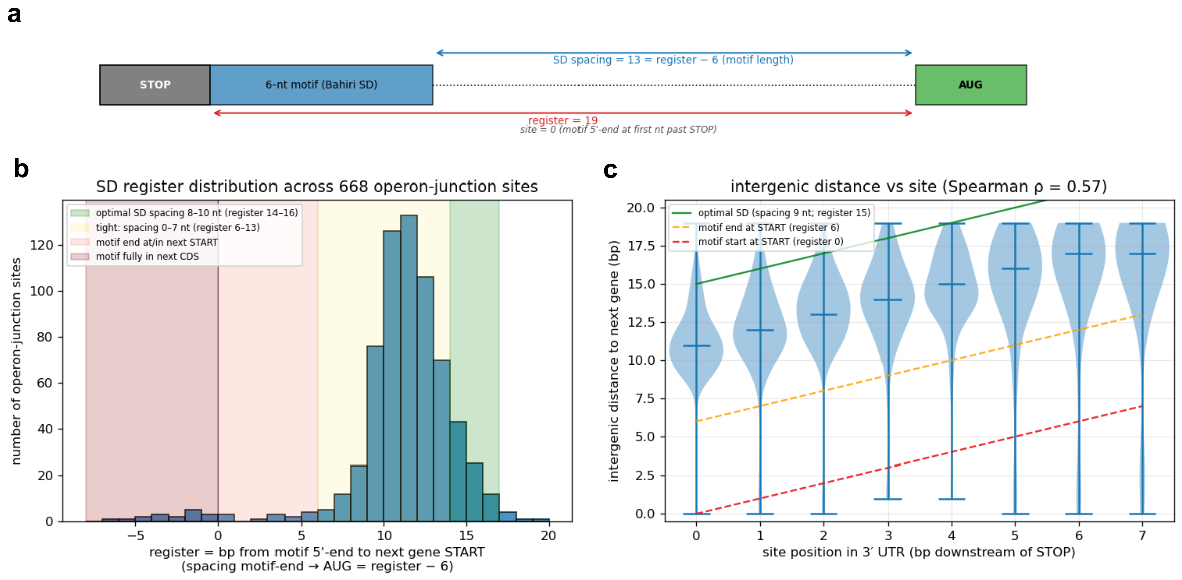


*Figure S 3:* ***Shine–Dalgarno register distribution at operon junctions.*** *(A) Schematic definition of the SD register relative to the upstream stop codon and downstream start codon.*
*(B) Distribution of SD register positions across 668 operon-junction sites. Shaded regions indicate optimal SD spacing, broader canonical SD spacing, motif overlap with the downstream start codon, and motif positions fully within the downstream CDS. Most sites cluster within the canonical SD spacing range. (C) Relationship between site position within the 3′ UTR and intergenic distance to the downstream gene. Dashed lines indicate expected intergenic distances for canonical SD spacing constraints. A positive correlation was observed between motif position and intergenic distance (Spearman ρ = 0.57).*

**Heatmap – 0% formaldehyde (no cross-linking).**
In the absence of formaldehyde cross-linking, and without an additional translation arrest step beyond rapid freezing in liquid nitrogen, ribosomal subunits are less stably associated with mRNA during the experimental procedure. Consequently, fewer subunit-associated footprints are recovered, as reflected in the reduced signal observed in the heatmaps (Figure S 4). In contrast, monosomes remain tightly bound to mRNA and therefore retain their footprints throughout the protocol, resulting in a detectable enrichment signal. Overall, the absence of cross-linking substantially reduces footprints originating from transient ribosomal subunit–mRNA interactions.

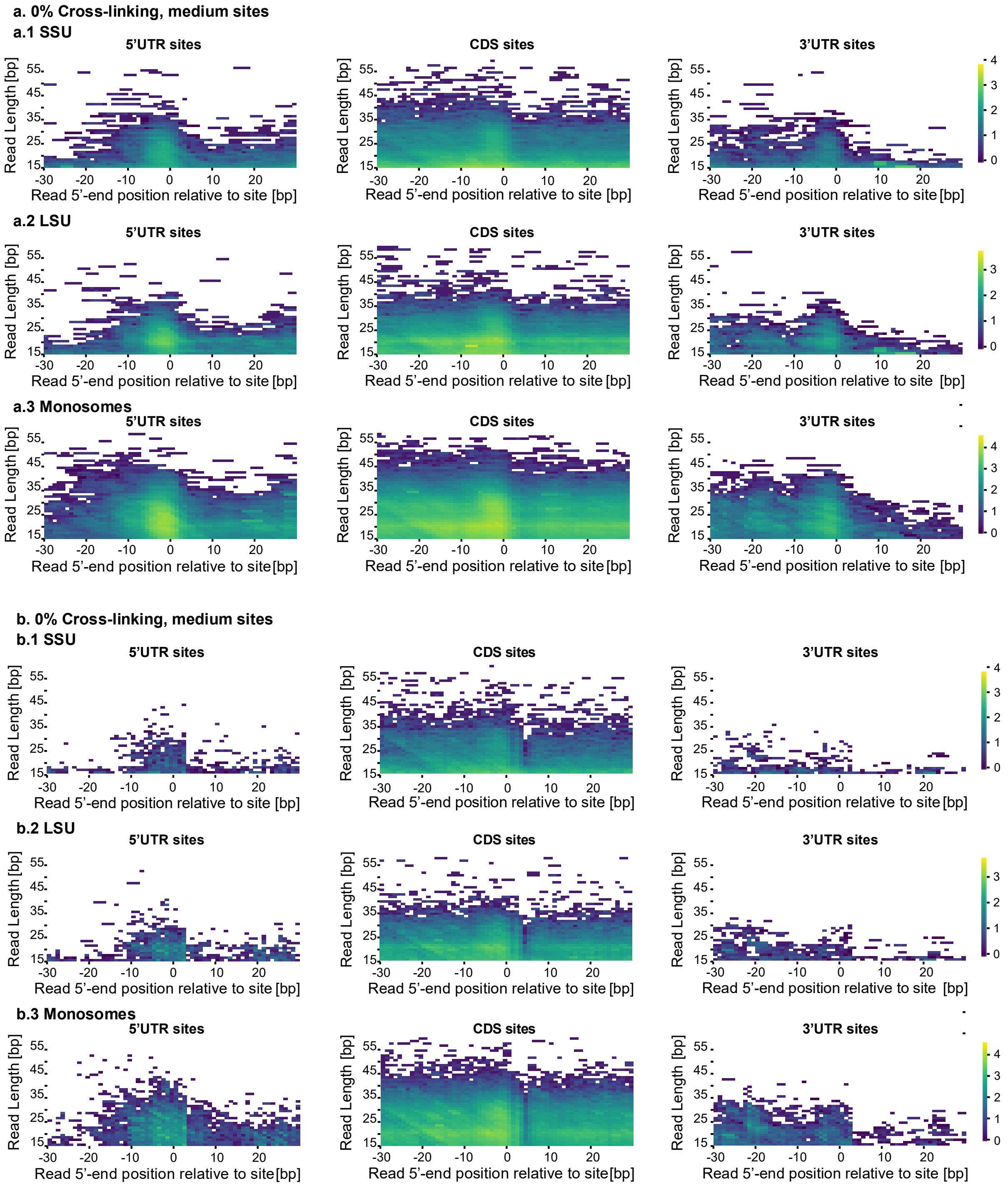


*Figure S 4:* ***Heatmap analysis of ribosomal subunit and monosome binding patterns across mRNA transcripts obtained by SSU-seq using 0.0% formaldehyde (no cross-linking)****. (a) Heatmaps corresponding to strong rRNA–mRNA hybridization sites. (b) Heatmaps corresponding to intermediate rRNA–mRNA hybridization sites. Heatmaps illustrate the distribution of ribosome-protected mRNA fragments across transcript regions: the 5′ untranslated region (5′ UTR, left), coding sequence (CDS, middle), and 3′ untranslated region (3′ UTR, right). Each row represents a different ribosomal species: (1) small subunit (SSU), (2) large subunit (LSU), and (3) monosomes. Color intensity (yellow) reflects the relative frequency of fragment mapping at each position.*

**Analysis of operon-junction-associated SSU enrichment in the 3′ UTR**

Strong hybridization sites within the first 8 bp of the 3′ UTR were stratified into operon-junction sites ($0 \le d < 20$ bp, n = 668) and non-junction sites ($d \ge 20$ bp, n = 761) based on signed intergenic distance to the next downstream same-strand gene. Operon-junction sites carried substantially higher SSU signal at the Shine–Dalgarno register than non-junction sites, with a raw mean ratio of 20.2-fold. To control for differences in gene expression between cohorts (gene-level CDS RPKM medians 22.9 versus 19.6; KS D = 0.105, p = 0.09), per-site signals were normalized by gene CDS RPKM. After normalization, operon-junction sites retained an approximately 7-fold higher signal (mean ratio 6.95×; one-sided Mann–Whitney U, $p < 1 \times 10^{-47}$).

SSU occupancy decreased monotonically with increasing intergenic distance (Spearman $\rho = -0.36$, $p < 1 \times 10^{-44}$ across positive distances) and remained concentrated within the canonical Shine–Dalgarno spacing range. Similar enrichment patterns were observed across alternative normalization procedures and read-length bands within the SSU-dominant range. Additional negative controls, including distant non-junction sites ($d \ge 100$ bp), CDS strong sites, and overlapping downstream genes ($d < 0$), all displayed near-baseline occupancy levels (Figure S 5).

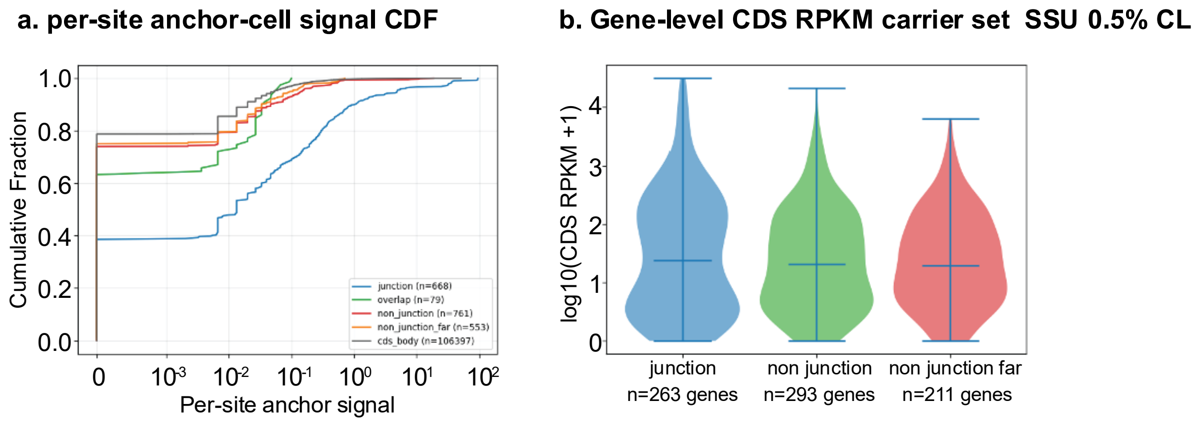


*Figure S 5:* ***Extended analysis of SSU occupancy at 3′ UTR strong hybridization sites.***
*Left: Cumulative distribution function (CDF) of per-site SSU occupancy across operon-junction, overlap, non-junction, distant non-junction, and CDS-body control site classes. Operon-junction sites display substantially elevated SSU occupancy relative to all control groups.Right: Gene-level CDS RPKM distributions for genes carrying operon-junction, non-junction, and distant non-junction sites, demonstrating comparable transcript abundance between cohorts.*

**0% formaldehyde (no cross-linking).**

To assess the contribution of cross-linking to footprint recovery, we analyzed samples generated without formaldehyde (0.0%) using the same metagene analysis applied to the 0.5% cross-linked samples (Figure S 6). In the absence of cross-linking, footprint enrichment near predicted hybridization sites was reduced and broader compared to the 0.5% samples. Nevertheless, overall trends were preserved, including SSU enrichment near predicted sites and continuous CDS coverage for monosome-associated footprints.

In contrast, 0.5% cross-linked samples displayed sharper and more pronounced enrichment peaks, particularly for SSU-associated footprints at strong hybridization sites. Differences between strong

and intermediate sites were also more clearly resolved in cross-linked samples. Monosome-associated footprints showed similar CDS coverage in both conditions, consistent with elongating ribosomes that remain tightly bound to mRNA.
These results indicate that formaldehyde cross-linking enhances recovery of transient SSU–mRNA interactions while preserving underlying ribosome distribution patterns.

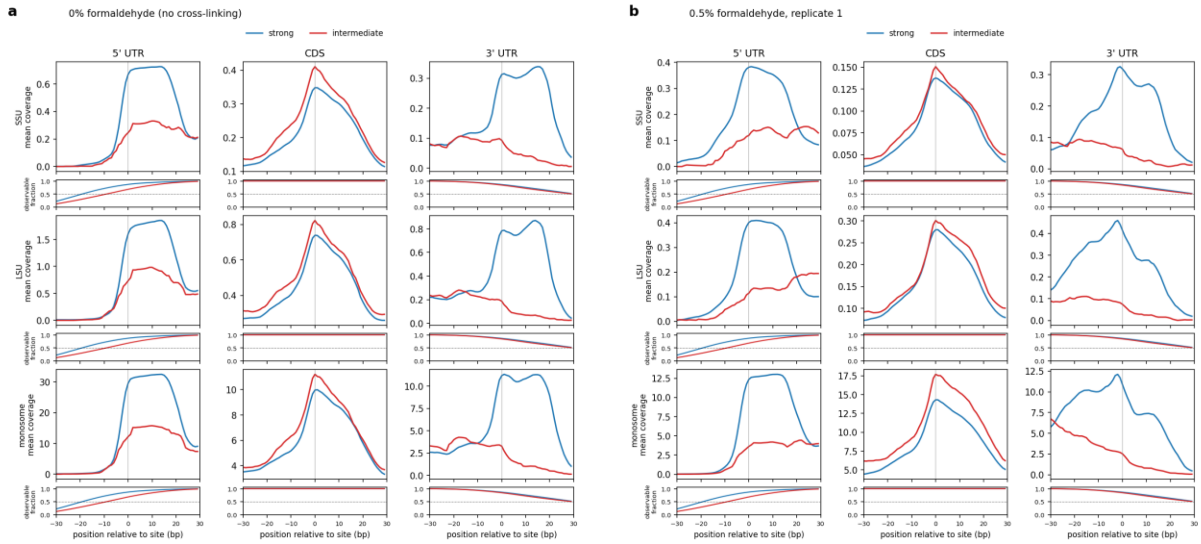


*Figure S 6:* ***Mean ribosome-protected footprint coverage around predicted rRNA–mRNA hybridization sites across mRNA regions for (a) a sample generated without formaldehyde cross-linking (0.0%) and (b) the first 0.5% formaldehyde biological replicate.*** *Profiles are centered on strong (blue) and intermediate (red) sites within a ±30 bp window; position 0 corresponds to the first nucleotide of each site. Rows correspond to SSU-, LSU-, and monosome-associated footprints (top to bottom), and columns to the 5′ UTR, CDS, and 3′ UTR. Coverage at each displayed position is normalized by the number of (read length, 5′-end position) combinations that could place a read there (see Methods); the strip beneath each profile shows the fraction of combinations possible at that position. The numbers of strong and intermediate sites are 14,477 and 1,618 in the 5′ UTR, 119,503 and 46,315 in the CDS, and 7,096 and 1,641 in the 3′ UTR. The libraries differ greatly in size (SSU libraries: 113,321 reads without cross-linking, 28,191 in replicate 1, and 3,430,376 in the replicate shown in Figure 5), so the y-axes are not comparable between panels or with Figure 5. This figure is presented for comparison with the 0.5% cross-linked sample shown in Figure 5 (replicate 2). Strong sites lie above intermediate sites in the UTRs and the profile shapes are similar to those in Figure 5, but fold enrichment over the CDS is lower in replicate 1 (for example, 2.8-fold for SSU at strong 5′ UTR sites, against 10.8-fold in Figure 5).*

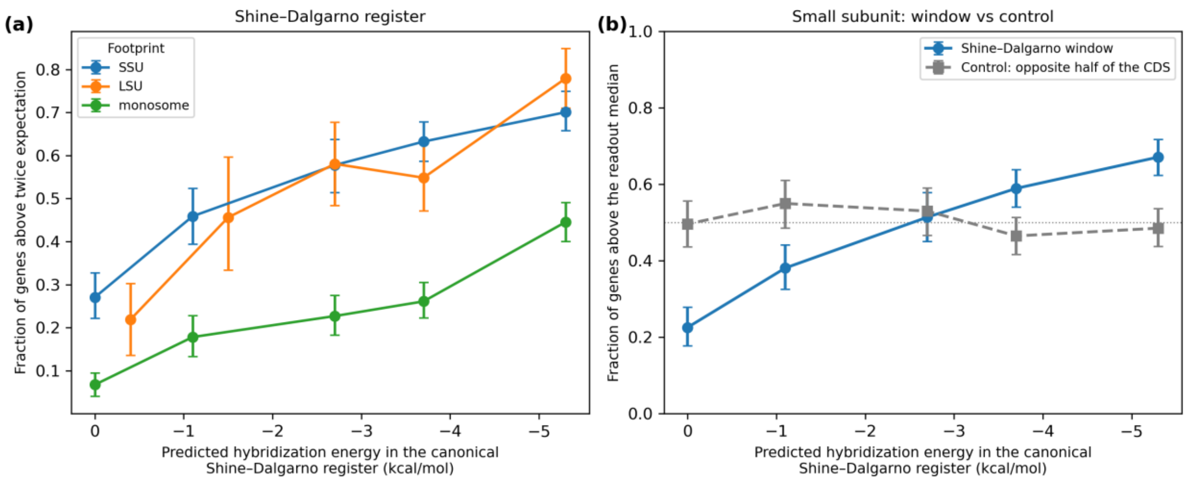


*Figure S 7*: Small subunit occupancy at the Shine–Dalgarno register scales with predicted hybridization energy at a fixed distance from the start codon. (a) Fraction of genes whose occupancy in a 16-nt window 15–30 nt upstream of the start codon exceeded twice its own expectation, against per-gene predicted hybridization energy in the canonical Shine–Dalgarno register, binned into quintiles; occupancy is normalized by the density in the first half of the same gene's coding sequence. (b) For the small subunit, the same readout compared against a control window in the opposite half of the coding sequence. Both readouts are binarized at their own medians, so that the control's null is as detectable as the effect it is testing; at the fixed threshold used in (a) the control's baseline is far lower and the comparison would be underpowered. The control is flat (Spearman ρ = +0.03, not significant); the large subunit and monosome fractions behave the same (ρ = +0.05 and +0.01, both not significant). Error bars are 95% bootstrap intervals over genes. Panels (a) and (b) use different binarizations, so their y axes are not comparable.

### Protocol: SSU-seq for *E.coli*

#### Reagent recipes

##### Section B – Cross-linkning:

- **Formaldehyde solution pH=8, 10ml**
  5ml Formaldehyde 37%
  5 ml PBS
  Add 1M NaOH until pH=8 is reached (about 10µl)
  Storage: prepare before use the exact amount, do not store the solution

- **Lysis buffer 10x 50 ml:**
  200 mM Tris pH 8
  1.5 M MgCl2
  1 M NH4Cl
  50 mM CaCl2
  4% Triton X-100
  1% NP-40
  Storage: -20C

##### Section D – Lysate extraction:

- **Low density buffer (50ml)**
  20mM Tris pH 8
  10mM MgCl2
  500mM NH4Cl
  2mM DTT

- **Resuspension Buffer (50ml):**
  20 mM Tris pH 8
  15 mM MgCl2
  100 mM NH4Cl
  5 mM CaCl2

##### Section F – Sucrose gradient

- **Sucrose buffer 10% (50%), 50ml: (**Prepare the gradients 1–3 h before use)
  20 mM Tris pH 8
  15 mM MgCl2
  0.1  M NH4Cl
  10% (50%)
  2mM DTT
  Store it at 4 °C for up to 1 week or at –20 °C for up to 6 months. Once DTT has been added, excess buffers should not be reused.

  **Section I – mRNA isolation**
  **Stop solution:**
  20% SDS
  0.4M EDTA (was dissolved in pH 8)
  DEPC water

#### Protocol

Note: All concentrations listed in recepis are final concentrations

##### A. Growth of bacterial cultures

1. Inoculate an overnight starter culture in MOPS medium 1ml (1:100ml starter:medium volume for mid-log growth) from a single colony.
2. Mid-log growth: The next day, prepare 250ml MOPS medium and add 2.5ml form OVN starter. Total of two 250ml beakers for each sample (total of 500ml).
3. Check OD after 3hrs and then once every ~half an hour. Return the culture immediately to 37°C after OD measurement
4. Stop the growth when reached to 0.5-0.6 OD, keep the culture on ice

##### B. Cross linking - Formaldehyde and Harvest by whole culture freeze

1. Weight the sorvalls prior to use

For one beaker of 250ml of E.coli mid-log culture:

2. Add *formaldehyde solution (pH=8)* (18% formaldehyde), for 0%, 1.0%, 1.5% concentration add 0ml, 13.5ml and 20.25ml respectively.
3. Mix for 5 minutes in RT (incubation)
4. Add 6.25ml of 2.5M glycine
5. Mix for 2 minutes in RT
6. While the reaction is quenching transfer the solution to tubes of 250ml, calibrate the weight before centrifuge. Fill the sorvalls only 2/3 of the volume.
7. Centrifuge the cells for 10 minutes, 4000rpm at 4C
8. Discard the supernatant to a Formaldehyde waste
9. Wash each pellet with about 40 ml of cold PBS and unite the pellets from of the whole sample (out of 500ml mid-log culture in total)
10. Centrifuge the cells for 10 minutes, 4000rpm at 4C
11. Discard the sup
12. Weight the pellet. Keep it on ice
13. Weight 3 tubes of 50ml
14. Wash the pellet again with about 15ml of cold PBS and transfer the cells to 50ml tubes
15. Centrifuge the cells for 5 minutes, 3200rpm at 4C. Discard the supernatant
16. Weight the pellet. Keep it on ice
17. Resuspend the pellet with 1ml MOPS
18. Drip into $\iota N_2$ the 1ml resuspended pellet together with 1ml of *lysis buffer* of 2X. Keep it frozen in $\iota N_2$ until pulverization.
    **Caution! Ensure that the liquid nitrogen is completely evaporated before sealing. Leftover liquid nitrogen in a sealed tube can cause a hazardous pressure build-up.**
    **PAUSE POINT - The frozen cells can be stored at –80 °C or below for at least 6 months.**

##### C. Cell lysis by freezer mill

The cylinder is good for about 4 grams. 1 batch of grinding:

1. Before the beginning of the first cycle: precool the Mixer mill for 5min
2. Transfer 1 frozen batch into the grinding cylinder
3. Grind using the following program: 5 cycles, 20Hz, 2min run, 1min cool, 5 min pre-cool
4. Transfer the pulverized cells into a pre-chilled 50 ml conical tubes contains $\iota N_2$
5. Store at -80 °C for later use or continue.
   Before grinding different CL reactions clean the mixer mill and then start section (C) in this protocol from the beginning.

6. Clean the mixer mill and its components. Rinse with DW and then with UPW and dry it with Methanol on a paper. Use the machine's parts when they are dry.

**PAUSE POINT - The frozen cells can be stored at –80 °C or below for at least 6 months.**

#### *D.* Lysate Extraction

1. Thaw tubes with pulverized bacteria in bain-marie with water of 30°C, thaw about 3 min, swirl gently occasionally during thawing.
2. As soon as the last pieces of the lysate thaw, transfer the tubes to ice, and then add 3.6 µl of RNaseOUT (40U per 100 mg of wet cell mass).
3. Vortex gently a further 1-2sec and then immediately put the tubes on the ice. **Perform all the subsequent operations on crushed ice unless otherwise indicated.**
4. Pellet cell debris by centrifugation of the tubes at 3200rpm for 10 min at 4 °C.
5. Carefully transfer the supernatants to a new 1.5-ml microcentrifuge tube. Avoid disturbing the pellets.
6. Pellet cell debris by centrifugation of the tube at 13,500g for 10 min at 4 °C.
7. Repeat Step 6. Document the volume collected.
8. Check that lysates are buoyant in 10% (wt/vol) sucrose before proceeding, by gently pipetting a small aliquot (~2 µl) of lysate into a spare tube of 10% (wt/vol) sucrose. If the lysate sinks, dilute it with low density buffer until it is buoyant.
9. Measure A260 of a 1:100 dilution using RNA suitable water
   Note: Calculate the amount needed for 12.5 AU (roughly 0.5 mg RNA), where AU = 1 ml solution with a given A260.
   If a sample is less than 12.5 AU, higher volume of sample is needed. As little as 5 AU can be used for library preparation, but this is not recommended.
   a. 12.5 AU/(A260 * 100/1,000) = µl needed (this takes into account the 100-fold dilution)
   b. Use the calculation above to calculate the volume needed for 12.5 AU, and add *Resuspension Buffer* to bring volume to 200 µl. the calculation is:
   X[AU]*100[dilution factor]*200[ul]/1000[ul]=y[AU]
   Where x is the result of the OD measurement, y is the absorbance in 200[ul] sample (amount of RNA in 200ul). Y needs to be compared to the 12.5[AU], amount that is recommended for library preparation. (The amount of Y [AU] there are in 200ul in comparison to 12.5 [AU])
   **PAUSE POINT - The frozen cells can be stored at –80 °C or below for at least 6 months.**

#### *E.* Nuclease digestion

Use 750U of MNase to digest 12.5 AU of sample in 200 µl. Note: If sample is less than 12.5 AU, scale the MNase accordingly.

1. Add 6 µl of SUPERase-In
2. Add calculated amount of MNase
3. Incubate at 25 °C for 1h shaking
4. Add 2 µl of 0.5 M EGTA pH 8 to quench the reaction (EGTA chelates calcium to quench MNase, a calcium dependent enzyme).
5. Return to ice until next step.

#### *F.* Sucrose gradient

**<u>Preparing tubes with light and heavy solutions</u>**

1. Preparing solutions 10% and 50% of sucrose
2. Mark the halfway point on the tube using the SW-41 marker-block
3. Draw the light solution of 10% sucrose with 60ml syringe and a cannula.
4. Exclude air from the cannula by up and down the solution
5. Add the solution in the tube slightly above the marked halfway point

6. Repeat for each tube with the same solution
7. Add the heavy solution (50% sucrose) using a 10ml syringe and a layering cannula:
   7.1. Exclude air from the cannula by up and down the solution
   7.2. Clean the outside of the cannula with paper towel
   7.3. Push a small droplet of liquid out of the end of the syringe, remove the droplet while ensuring there is no air in the cannula
   7.4. Insert the cannula to the bottom of the tube
   7.5. Gently add the solution without mixing between the light and the heavy solutions
   7.6. Keep the canula below the interphase of the solutions
   7.7. Fill the tube until the heavy solution reach the marked halfway point
   7.8. Make sure to leave empty 2mm in the tube
   7.9. Withdraw the cannula from the side of the tube and straight up
   7.10. Close the tube with a rubber cap, the hole on the cap is the last to seal the tube
8. Place the tubes in the tube holder rack
9. Create the gradient in the BIOCOMP fractionator

#### G. Ultracentrifuge of the gradient with samples

1. Clean the buckets of rotor SW-41 before use and let them dry over night
2. Remove the cap from the tube starting with the hole (the hole is where the protrusion in the cap)
3. Wipe the tube with the sucrose gradient and insert into the bucket (document the number of the bucket and the name of the sample)
4. Remove from the top of the tubes' sucrose gradient 200µl (same as the samples' volume), using protected tips. Keep the removed solution until the weight of the bucket is balanced and equivalent to its pair
   --> The difference in weight in Pair of buckets that face each other in the ultracentrifuge, is maximum 10mg
5. Add the sample on top of the gradient solution in the tube, 200µl
6. Close the buckets with its cover using screwdriver
7. If additional volume is needed for balancing use 10% sucrose solution
8. If there are empty buckets they must be balanced as well. It is possible to balance with tube filled with water

The buckets are ready to be placed in the ultracentrifuge

9. Use Rotor SW-41, volume (according to the software) 12.5ml open top
10. Place the buckets on the rotor – place the buckets according to their numbers
11. Set the ultracentrifuge for 35Krpm (201,000g), 4ºC, 3.5hr and use acceleration and deceleration setting of maximum speed
12. After the ultracentrifuge cycle is done: take the buckets gently, preferred on ice, to the gradient station

#### H. Creating profile in the gradient station

The protocol in details is described at the end of this protocol.

1. Prepare sucrose gradient fraction collector following the manufacturer's instructions.
2. Zero the gradient station
3. Create the polysome profile
4. Divide each collected fraction to batches of 350µl
5. Freeze fractions in liquid nitrogen and store at -80 °C

**PAUSE POINT - The frozen fractions can be stored at –80 °C or below for at least 6 months.**

#### *I.* mRNA Isolation - Phenol chloroform extraction

1. Thaw the relevant fraction
2. Add stop solution to 10% (vol/vol) (40 µl 100%), Tris–HCl (pH 7.4 at 25 °C) to 10 mM (4 µl, 1 M), glycine to 10 mM (1.6 µl, 2.5 M), Rnaseout and deionized nuclease-free water (4.4 µl if the starting fraction volume was 350 µl) to obtain 400 µl as the final volume.
3. Vortex and spin-down.
4. Place the tubes with the mixtures into a thermomixer and continue shaking at high speed (1,300 rpm) for 20 min at 65 °C to reverse the cross-links. During that period open the tube to release pressure
5. Chill on ice for 5min
6. Add Trizol and chloroform according the steps describes in manufacturer's instructions
7. Ethanol precipitation

**Elaboration to section H**

**Creating the gradient**

Make sure the instrument is leveled:

1. place the bubble level from back to front on the gradient station – change the height of the stations' legs for leveling
2. Turn on the gradient station
3. Press "GMST" to level the magnetic plate (the right wing of the gradient station)
4. Place the bubble level horizontally on the magnetic plate and press "up" or "down" until it's leveled.
5. Press "Done"
6. Place the tube holder rack with the tubes on the magnetic plate
7. Press "Grad", choose from the following options:
    1. List – all programs
    2. Last – the last program used in the lab
    3. Recent – all gradients used in the lab
8. Choose the relevant program: SW41 rotor, gradient: 10%-50% sucrose
9. Press "use"
10. Press "Run"

The magnetic plate with the tubes on top of it, starts to move to create the gradient.

**Polysome profiling**

**Part 1: zero the gradient station**

1. Turn On the gradient station
2. Rinse for 10 sec – use the toggle valve on the right hand of the gradient station in the up position
3. Follow step 1 in the cleaning protocol
4. Connect the computer, Triax software, to the gradient station:
    1. Press "Scan"
    2. Make sure 2 USB cables are connected to the computer, the other ends are the triax and the gradient station
    3. Open the software in the computer

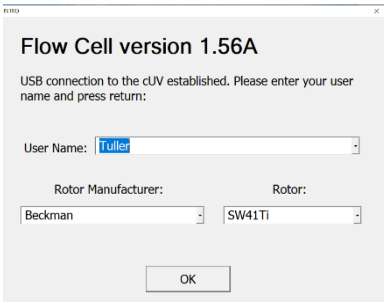


4. Fill in the software: Username, name of experiment, other details and press the “LED power”

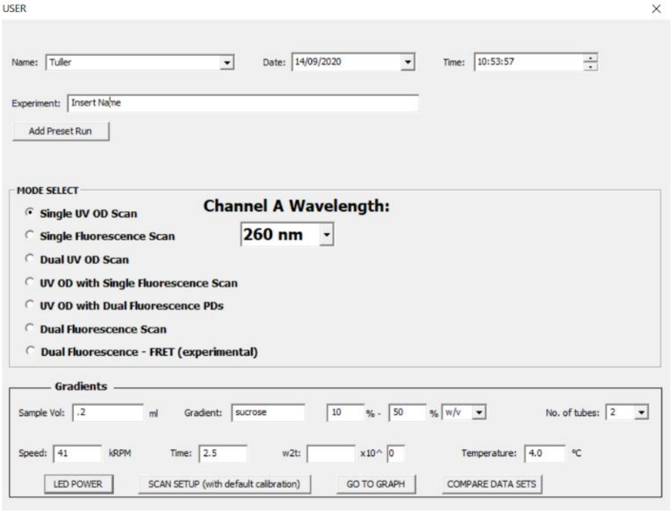


To get the source and sample values into the optimal range
source >100 000
sample >800 000 , <900 000
you need to adjust the ontime on the LED power page.

5. Let it run while rinsing the gradient station – the numbers in the software should be around 80K. If the numbers are low rinse again.
6. Press the “scan setup” – make sure to choose the “slow start point” and absorbance of 260nm
7. Zero the gradient station – press “zero” in the software. A stable profile around 0 should be formed in the chart. After the chart is stable the gradient station is ready for use.
8. If the chart is not around 0, perform step 2 and 3 again and press “another zero”. Check if the chart is stable around the zero (in y axis).

**Part 2: Creating a profile**

1. Take off the rinse adaptor and screw in the piston adaptor (black nozzle with a transparent silicone cap)
2. Take out the tube with the gradient solution and the sample, out of the bucket using tweezers
3. On top of the tube place the cap of the tube holder

4. Place the tube with the cap inside the tube holder and lock the cap inside, make sure it’s locked
5. Place the tube holder on the gradient station; above the lamp and below the piston.
6. Put the tube holder on top of the lamp, while the transparent panel is facing to the right
7. Lock the tube holder in its place by turning it 90°to the left, so the transparent panel is facing forward.
8. Press “start” in the software
9. Press “air” on the gradient station for 10sec
10. Press “OK”

**The profile is formed, collect the relevant fraction**

14. After the profile is created, save the profile
15. Take out the tube holder and the tube out.
16. Clean before placing a new tube sample - Follow step 1 in the cleaning protocol

Start from Part 2 step 6 to create another profile

#### Analysis of sequencing reads

Proper trimming of the reads is important for efficient mapping. Here we provide some guidance on the use of (1) cutadapt (Martin, 2011) to remove LACE-seq 3’linker, (2) UMI-tools `extract` (Smith, 2017) to move the UMI sequence from the read to the read name so that PCR duplicates can be removed after the alignment, (3) cutadapt to remove the T preceding the RPF.

Trimming is done in 3 steps:

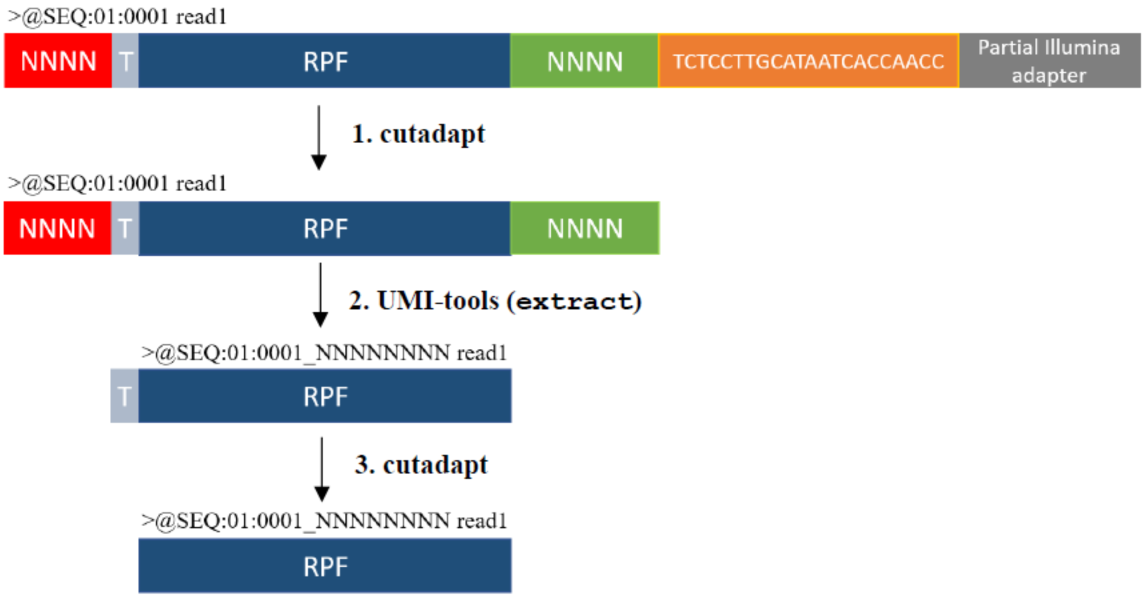

**STEP1: cutadapt**

First the LACE-seq linker is trimmed from the 3' end of each read and only reads longer than **X+9** nt are retained, while shorter reads are discarded:

```
cutadapt --minimum-length X+9 -a TCTCCTTGCATAATCACCAACC --discard-untrimmed
-o trim.fastq input.fastq
```

| Parameter | Definition |
|---|---|
| --minimum-length X+9 | Reads are retained if they are longer than X+9 nt, where X is the length of the RPF (usually X=20 for ribosome profiling analysis), and 9 is the sum of the lengths of the 5' and 3' UMIs |
| -a TCTCCTTGCATAATCACCAACC | Removal of the LACE-seq 3' linker and any sequence that may follow |
| --discard-untrimmed | Reads in which *no* adapter is found are discarded |
| -o trim.fastq | The output file name |
| input.fastq | The input file name |

**STEP2: UMI-tools (extract)**

The sequence of the 5' and 3' UMIs are moved from the read sequence to the read name:

```
umi_tools extract -I trim.fastq --bc-
pattern='^(?P<umi_1>.{4}).+(?P<umi_2>.{4})$' --extract-method=regex -S
extract.fq
```

| Parameter | Definition |
|---|---|
| -I trim.fastq | The input file name must be the same as the output file name in step1 |
| --bc-pattern='^(?P<umi_1>.{4}).+(?P<umi_2>.{4})$' | extract the first 4 (5'UMI) and the last 4 bases (3'UMI) of each read |
| -S extract.fq | The output file name |

NOTE: UMI-tools dedup can be used after alignment to remove duplicates based on the mapping coordinate and the UMI attached to the read name.

**STEP3: cutadapt**

The T preceding the RPF is then removed:

```
cutadapt -g ^T --discard-untrimmed -o trim2.fastq extract.fastq
```

| Parameter | Definition |
|---|---|
| -g ^T | Removal of the first T at the start of each read |
| -o trim2.fastq | The output file name |
| extract.fastq | The input file name must be the same as the output file name in step2 |
